# Decoding Smell from Receptor Structure

**DOI:** 10.64898/2026.04.22.720159

**Authors:** Hsiu-Yi Lu, Aashutosh Vihani, Maira H. Nagai, Xiaoyang S. Hu, Dan Takase, Conan Juan, Claire A. deMarch, Hiroaki Matsunami

**Affiliations:** Department of Molecular Genetics and Microbiology, Duke University School of Medicine, Research Drive, Durham, NC 27710 USA; Department of Neurobiology, Neurobiology graduate program, Duke University Medical Center, Durham, NC 27710, USA; Department of Neurobiology, Duke University School of Medicine, Research Drive, Durham, NC 27710 USA

## Abstract

Olfaction enables animals to detect and discriminate a vast array of chemicals, yet how odorant receptors (ORs) encode ligand selectivity remains unclear. Although recent advances in protein structure prediction have expanded access to OR structures, linking these to function at scale has been challenging. Here, we combined AlphaFold3-predicted receptor structures with protein language model embeddings and in vivo pS6-IP-Seq measurements of olfactory sensory neuron activation across a chemically diverse odor panel to train a deep learning model of OR-ligand interactions. The resulting framework predicts receptor responses, organizes receptors by functional similarity independent of sequence, and identifies structural determinants of ligand selectivity. These findings establish a structure-based map of OR function and provide a foundation for predictive and interpretable models of olfactory coding.

## Introduction

Animals rely on multiple sensory systems to navigate and interpret their environment, each specialized to encode distinct types of physical or chemical information. Among these, olfaction presents one of the least understood and most complex encoding challenges. In color vision, the task is to discriminate between wavelengths of light and is achieved through the tuning of a small number of cone opsins with distinct spectral sensitivities, which are a subfamily of class A G protein–coupled receptors (GPCRs), is central to how humans and other animals perceive light as distinct colors (*1, 2*). Odorant receptors (ORs) in vertebrates are also class A GPCRs; however, in contrast to vision, olfaction must distinguish among diverse chemical molecules, requiring large repertoires of odorant receptors (ORs). ORs ranging from hundreds to thousands to encode this high-dimensional chemical space, which constitute the largest subfamily of class A GPCRs (*3*), with humans expressing approximately 400 intact ORs and mice over 1,000 (*4, 5*). Mammals devote a substantial fraction of their genomes to ORs, which reflects the evolutionary importance of chemical sensing across species. Despite their central role in odor perception, a fundamental unresolved question is how an individual OR determines which chemicals it binds and how it responds across chemical space. Addressing this question is essential for understanding the logic by which chemical information is encoded at the receptor level, yet progress has been limited by long-standing technical barriers to obtaining high-resolution OR structures.

Recent cryo-EM studies have resolved structures of mammalian ORs, first with OR51E2 in complex with propionate (*6–12*). Notably, these cryo-EM structures revealed that odorants primarily occupy a conserved binding cavity location, suggesting a canonical structural locus for odorant recognition across ORs. Functional studies further demonstrate that mutations altering the shape or chemical properties of this cavity can shift ligand preference, suggesting ligand selectivity is primarily influenced by the geometry and physicochemical features of the binding pocket.

In parallel, advances in computational biology have transformed access to protein structural and evolutionary information. Deep learning-based structure prediction methods such as AlphaFold3 (*13–15*), have provided the potential for accurate *in silico* predictions for the entire OR repertoire, offering a complementary resource where experimental structures are lacking. Additionally, other approaches like protein large language models (LLMs), such as the Evolutionary Scale Model (ESM) (*16*) extend this progress by embedding sequence-based evolutionary information into learned vector spaces that infer functional and biophysical constraints (Lin et al., 2023). Together, these approaches overcome a major limitation by providing receptor-level structural information at scale, creating an opportunity to pursue structure-informed modeling of OR–ligand interactions.

Many existing efforts to model olfactory perception and OR–ligand interactions have focused on ligand descriptors, psychophysical measurements, and receptor sequence features, yielding important insights into odor coding and chemical space (*17–21*). These approaches offer interpretable representations of odorant chemistry and perceptual outcomes. However, the limited availability of high-resolution receptor structures has limited the ability to connect ligand binding to three-dimensional features of the receptor binding cavity. Given that the receptor structure plays a central role in determining ligand specificity and activation (*6, 8*), integrating structural information is essential for directly linking chemical recognition to receptor function.

To address this gap, we developed an end-to-end modeling framework that integrates AlphaFold-predicted OR structures with ESM2-derived sequence embeddings within convolutional neural networks to predict OR–ligand interactions. Central to this effort is the use of pS6-IP-Seq, an in vivo assay that captures activation of olfactory sensory neurons through enrichment of phosphorylated ribosomal protein S6–associated transcripts (*22, 23*). While numerous studies have reported OR–ligand interactions, these datasets are typically small, heterogeneous in experimental design, and difficult to integrate across studies. In contrast, collection of pS6-IP-Seq studies enables systematic, parallel measurement of receptor activation across chemically diverse odorants within a unified experimental framework, making it particularly well suited for machine learning applications.

By integrating large-scale pS6-IP-Seq activation data with predicted receptor structures and sequence-derived embeddings, we set out to ask whether receptor structure representation can be explicitly leveraged to model and interpret OR–ligand interactions. Specifically, we aimed to determine the structural features of the receptor that encodes for the ligand selectivity and receptor tuning. Through this approach, we seek to establish a structure-informed framework for linking receptor feature to odorant activation, and to assess whether focusing on a canonical binding cavity can provide a mechanistic basis for understanding olfactory coding at the receptor level.

## Results

### pS6-IP-Seq as a Strategy to Identify Odorant Activated OR In Vivo

The phosphorylation of the ribosomal protein S6 is a well-established marker of neuronal activity (*24*). Similar to the induction of immediate early genes such as c-Fos, pS6 reflects recent neuronal activation, but with the added advantage of being physically associated with mRNA molecules. As a result, phosphorylated ribosomal S6 (pS6) complex can be immunoprecipitated and coupled to RNA sequencing, enabling a direct molecular readout of transcriptional profiles from activated olfactory sensory neurons (OSNs) following odor exposure (Fig. 1A). Given, each mature OSN expresses a single olfactory receptor gene (*25–29*), we can directly assign OR identities to individual OSNs. This organization allows pS6-IP-Seq to link odor-evoked neuronal activity to specific OR identities in vivo. Consistent with this framework, previous studies have demonstrated the utility of pS6-IP-Seq in inferring odorant–OR interactions in vivo freely behaving mice (*22, 30, 31*).

**Figure 1.**
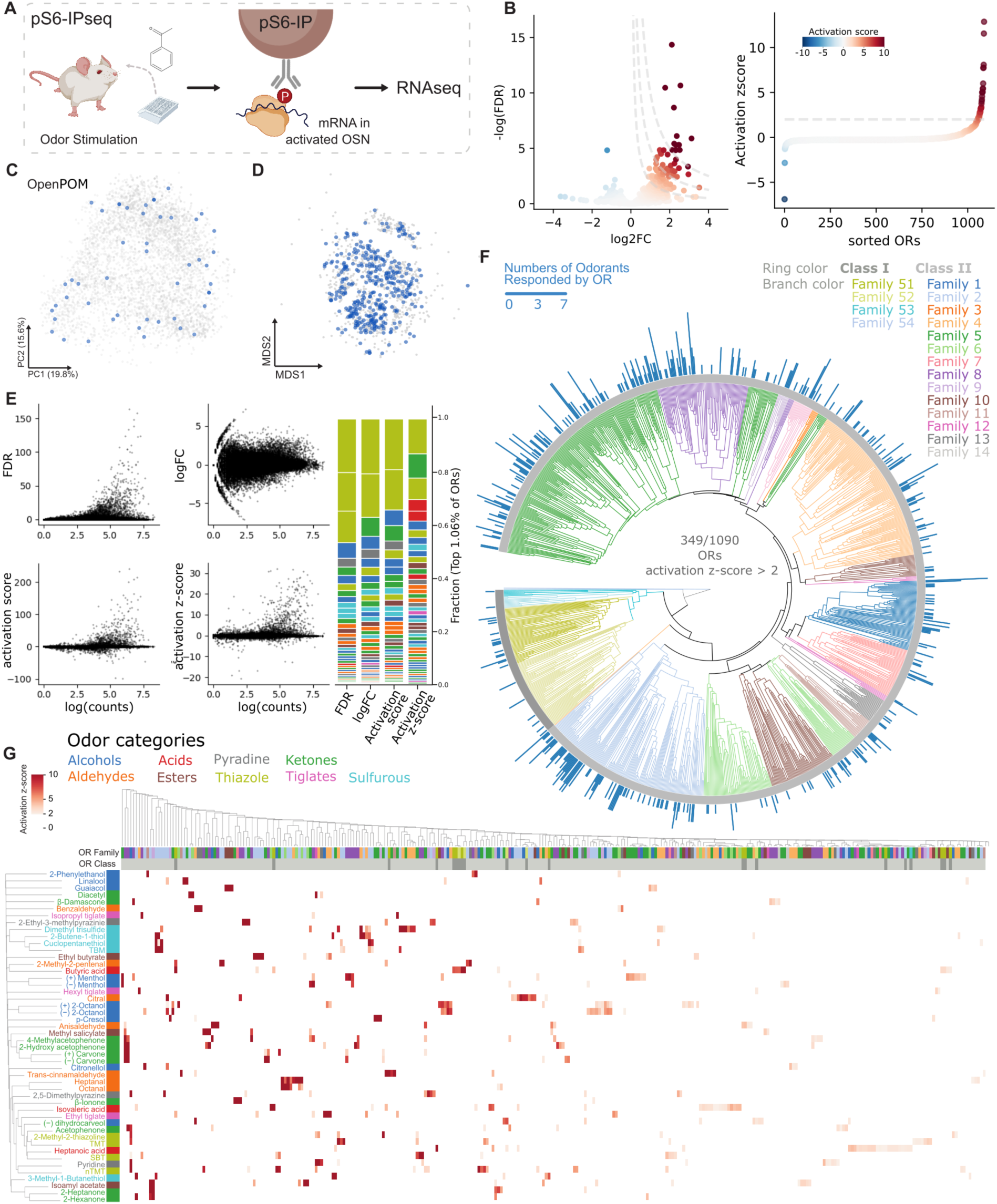
Defining odor-evoked olfactory receptor activation using pS6-IP. **A**. Schematic of the pS6-IP-Seq experimental workflow. Odor exposure induces phosphorylation of ribosomal protein S6 in activated olfactory sensory neurons (OSNs). pS6-IP-Seq enriches mRNAs expressed in activated OSNs, which are subsequently profiled by RNA sequencing to identify odor-responsive olfactory receptors (ORs). **B**. Quantification of OR activation in response to 100% acetophenone stimulation. Left: Volcano plot showing enrichment of OR mRNAs, with points colored by activation score. Right: ORs ranked by activation z-score, with colors indicating activation score. The horizontal dashed line denotes the activation z-score threshold (> 2) used to define activated receptors. **C**. Chemical perceptual space representations of tested odorants. Principal component analysis (PCA) of 4680 small molecules based on OpenPOM. The 48 odorants tested by pS6-IP, each activating at least one OR, are highlighted in blue. **D**. ORs space representations of responding receptors. Multidimensional scaling (MDS) visualization of intact ORs based on pairwise Grantham distances. ORs responding to at least one of the 48 odorants are shown in blue (n = 349). **E**. Scatter plots of different metric’s (FDR, logFC, activation score, activation z-score) relationship biases with receptor counts. Right: Histogram showing the fraction of identified top 1% odor–ligand pairs selected using different response metrics. **F**. Circular phylogenetic tree of ORs, with branch colors indicating OR family and shaded ranges denoting OR class. The outer ring displays a bar plot representing the number of odorants to which each OR responds, as defined by activation z-score threshold > 2. **G**. Heatmap summarizing activation z-scores across all 48 pS6-IP odor experiments with activation z-score threshold > 2. Rows represent odorants and columns represent ORs, with hierarchical clustering applied to both dimensions.

To ensure broad coverage of chemical space, we selected a panel of 48 odorants spanning diverse functional groups, molecular scaffolds, and physicochemical properties from panel of small molecules commonly found in foods and fragrances (*32*). In addition to maximizing global chemical diversity, the panel was intentionally designed to include closely related chemical pairs, such as enantiomers and homologous series differing by carbon chain length, enabling assessment of receptor sensitivity to subtle structural variations. Projection of these odorants into a reduced chemical feature space using Morgan fingerprint based chemical features (*33*), as well as clustering within an Open Principal Odor Map (OpenPOM) derived perceptual space (*21, 34*), generated from message passing neural networks trained on human odor perception, confirmed that the selected panel captures both structural and perceptual diversity across odorants (Fig. 1C and fig. S1B).

### Defining Activation Score to Quantify OR Responses

Traditional identification of activated olfactory receptors in pS6-IP experiments often repurposes differential expression criteria developed for gene-level analyses, such as false discovery rate (FDR) thresholds (e.g., FDR < 0.05) or minimum log fold-change (logFC > 1). While effective for detecting broadly regulated transcripts, these metrics are not optimized for classifying receptor activation and introduce systematic biases when applied to ORs. LogFC is strongly influenced by transcript abundance and sequencing depth: ORs with low counts can exhibit inflated fold changes due to small denominators, whereas highly expressed receptors frequently display more modest logFC values despite reproducible enrichment. As a result, logFC-based selection preferentially highlights sparsely expressed ORs while deprioritizing abundantly expressed receptors, confounding biological activation with sampling effects (Fig. 1E). In contrast, FDR-based selection introduces a complementary but distinct bias. Because FDR reflects statistical confidence rather than effect size, it preferentially prioritizes receptors with stable baseline expression and low variance, even when the magnitude of enrichment is small. Consequently, receptors exhibiting consistent but weak changes can be classified as activated, whereas biologically meaningful but variable responses may be excluded. Together, reliance on either logFC or FDR alone emphasizes different statistical regimes rather than receptor activation, limiting their interpretability for OR classification.

To address these limitations, we developed a composite metric, termed the Activation Score, which integrates both enrichment magnitude and statistical confidence without relying on a single criterion alone.

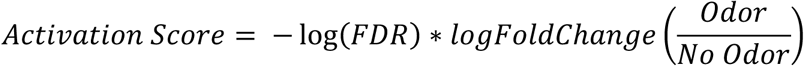

By jointly considering effect size and reliability, the Activation Score captures activation signals that reduces sensitivity to transcript abundance, variance structure, and arbitrary threshold selection. The advantage of this approach is illustrated in volcano plots of acetophenone pS6-IP responses (Fig. 1B), where increasing Activation Score thresholds yield a more balanced identification of responsive ORs compared with conventional FDR- or logFC-based filtering (Fig. 1E).

Furthermore, because different odorants elicit varying overall activation strengths at fixed experimental concentrations, direct comparison of raw activation scores across odors can introduce additional bias. To mitigate this, we normalized activation scores by z-scoring within each odor experiment, generating an odor-specific activation z-score. This normalization ensures that OR detection reflects relative responsiveness within an odor condition rather than absolute signal magnitude. Using this normalized metric, we defined activated ORs as those with an activation z-score greater than 2 for all downstream analyses.

Comparison of the top 1.06% of OR–odorant pairs, corresponding to the number of pairs exceeding an activation z-score of 2. Whereas FDR- and logFC-based criteria were skewed toward small subsets of odorants, activation z-score–based selection preserved a more diverse distribution of odor categories (Fig. 1E), indicating reduced bias toward ligands that produce exceptionally strong or abundant signals (*35*).

Analysis of OR responses identified using the activation z-score revealed broad coverage of receptor space. Projection of activated ORs into a sequence-based receptor space defined by Grantham distances demonstrated that activated receptors were widely distributed rather than concentrated within specific OR families or classes (Fig. 1D). This result was further supported by phylogenetic analysis, in which activated ORs were dispersed throughout the OR tree (Fig. 1F). Together, these analyses indicate that pS6-IP enables deorphanization across a diverse and representative fraction of the ORs.

Consistent with this diversity, clustering of OR–odorant responses across all 48 odor’s pS6-IPseq revealed that receptors did not group according to family identity, which is defined by overall amino acid similarities (*36*), but rather according to shared ligand responsiveness (Fig. 1G). Conversely, ORs with high sequence similarity did not necessarily respond to the same odorants, highlighting the dissociation between phylogenetic proximity and functional tuning.

Together, the odor-normalized Activation Score z-scored provide a more balanced and interpretable framework for quantifying OR activation from pS6-IP data. This metric reduces bias associated with transcript abundance and differential response strength, forming a robust foundation for the downstream structural modeling and machine learning analyses presented in this study.

### Encoding Odorant Receptor Structure

Having established a large-scale, chemically diverse dataset of in vivo OR activation, we next asked whether receptor structure contains information to predict ligand selectivity.

Specifically, we sought to determine whether ORs could be represented in a structure-informed space that reflects their likelihood to respond to a given chemical stimuli, thereby enabling an interpretable model of receptor coding.

We hypothesized that the three-dimensional architecture of the receptor, particularly features within the ligand-binding cavity encodes constraints that shapes chemical recognition. Under this hypothesis, structural representations alone should organize ORs into a space that reflects their patterns of chemical responsiveness, even without incorporating ligand features.

To establish a reliable structural foundation for this analysis, we first evaluated the ability of AlphaFold3 to generate accurate, activation-relevant OR conformations. As a test case, we modeled OR51E2 in complex with a mini Gαs protein using AlphaFold3 multimer. Notably, inclusion of the mini-Gαs protein alone was sufficient to stabilize an active-like receptor conformation, without providing the ligand (fig. S3). Comparison of the predicted structure with the recently solved cryo-EM structure of OR51E2 (*6*) revealed less than 1Å RMSD overall backbone deviation, indicating close agreement between predicted and experimental models (Fig. 2A and fig. S3). This result is consistent with prior observations that AlphaFold predictions tend toward inactive-like states when modeled in isolation but can be shifted toward active conformations through inclusion of signaling partners such as G proteins (Fig. 2A and B)(*6*).

**Figure 2.**
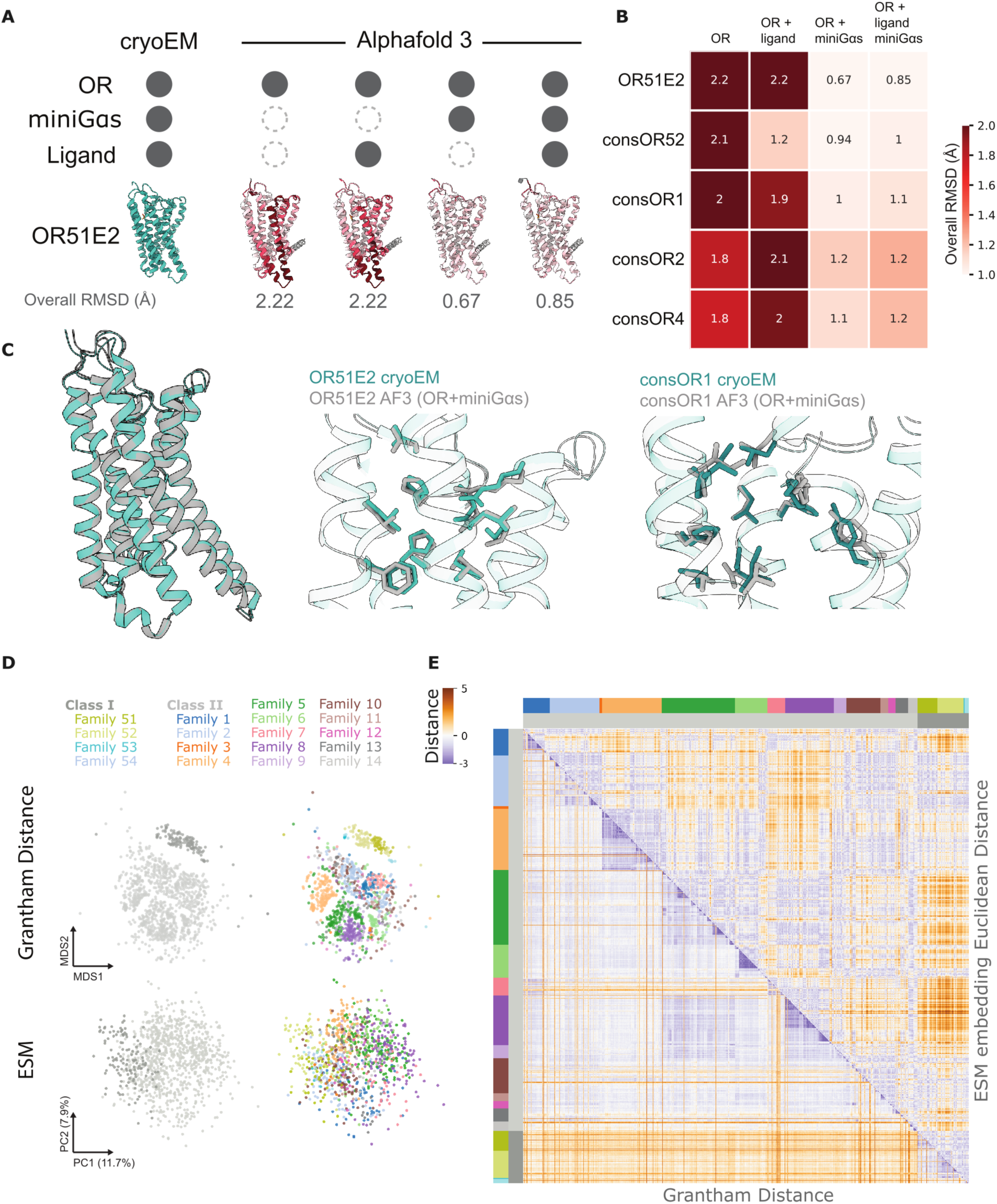
Structural accuracy of AlphaFold3 models and comparison of sequence- and language model–based receptor representations. **A**. Structural comparison between AlphaFold3-predicted models and the cryo-EM structure of OR51E2. Columns show different multimer configurations used for structure prediction. Structures are colored by per-residue RMSD relative to the cryo-EM reference, with overall RMSD values reported below each model. **B**. Heatmap summarizing overall RMSD values between AlphaFold3 predictions and experimentally determined cryo-EM structures across five available OR structures. **C**. Structural overlays of predicted and experimental OR structures. Left: superposition of OR51E2 cryo-EM and AlphaFold3 (OR+miniGαs) models. Middle: Magnified view of binding cavity residues for OR51E2, comparing cryo-EM and AlphaFold3 models. Right: Magnified view of binding cavity residues for consOR1, comparing cryo-EM and AlphaFold3 models. **D**. Low-dimensional visualization of OR similarity based on sequence-derived and language model–derived representations colored by class and family. Top: Multidimensional scaling (MDS) of pairwise Grantham distances. Bottom: Principal component analysis (PCA) of ESM-2 sequence embeddings. **E**. Pairwise distance heatmap showing Grantham distance based similarities (lower triangle) and Euclidean distances computed from ESM-2 embeddings (upper triangle) across ORs.

Importantly, similar structural concordance was observed across additional OR cryo-EM structures spanning distinct classes and subfamilies, with AlphaFold3 models showing consistently low backbone deviations. Together, these results support the use of AlphaFold3 with OR with miniGαs to generate active-like structural models for diverse ORs (Fig. 2B and fig. S3).

We next assessed whether these predicted structures were suitable for interrogating ligand selectivity by examining the architecture of the binding cavity Analysis of residues lining the predicted ligand-binding pocket showed that AlphaFold3 accurately recapitulates not only the overall receptor fold but also the positioning of key side chains that define cavity shape (Fig. 2C). The close correspondence between predicted and experimentally observed side-chain orientations indicates that AlphaFold3 models preserve the structural features critical for defining the binding pocket, supporting their use for downstream structural encoding and machine learning analyses.

### Feature encoding of primary sequence

To examine how sequence-derived representations capture OR similarity, we leveraged embeddings generated by the Evolutionary Scale Model (ESM), a protein language model trained on large-scale protein sequences to learn contextual and evolutionary patterns directly from sequence data (*16*). Unlike traditional representations based on primary amino acid identity or predefined substitution matrices, ESM embeddings encode each residue in the context of its surrounding sequence enabling the representation to capture higher-order relationships. ESM-based embeddings have been shown to correlate with protein structure, mutational effects, and functional similarity across diverse protein families (*37, 38*). To compare against a baseline measure of OR similarity from primary sequence, we computed pairwise distances using Grantham’s amino acid substitution matrix. These distances capture differences in physicochemical properties such as polarity, volume, and composition, and thus provide a familiar framework for comparing receptor families. Using multidimensional scaling, we projected the pairwise Grantham distances into two dimensions (Fig. 2D). The resulting clusters closely mirrored known OR class and family designations, confirming that physicochemical substitution metrics reflect evolutionary relationships across the OR repertoire as expected.

Full-length OR sequences were encoded using ESM, and the resulting embeddings were projected into principal component space for visualization (Fig. 2D). Compared with Grantham distance–based clustering, which primarily reflects amino acid similarity and substitution patterns, ESM embeddings showed weaker segregation by canonical OR families, suggesting that they capture sequence features beyond simple residue identity (Fig. 2D-E).

To directly compare these representations, we computed pairwise distances using both ESM embeddings and Grantham metrics. While the resulting distance matrices showed broad agreement at the level of OR class and family organization (Fig. 2E), they also revealed distinct patterns unique to each approach. Correlation analysis confirmed a moderate but incomplete relationship between the two measures (fig. S2), indicating that ESM embeddings encode additional evolutionary and contextual information not captured by traditional substitution-based metrics. Together, these results highlight that ESM provides a complementary, higher-order representation of OR sequence space.

### Integration of Structural Voxels and ESM Embeddings

With validated structural models and residue-level embeddings in hand, we developed a unified spatial learning framework that integrates three-dimensional receptor architecture with sequence-derived features into a single representation (Fig. 3A). For each OR, the cavity space was calculated using pyKVFinder (*39*) from the AlphaFold3-predicted structure generated in complex with mini-Gα. These cavities were then filtered against the canonical binding cavity reference, yielding a single, standardized ligand-binding region for each receptor. Amino acid residues encasing this cavity space were extracted as the structural representation of the OR.

**Figure 3.**
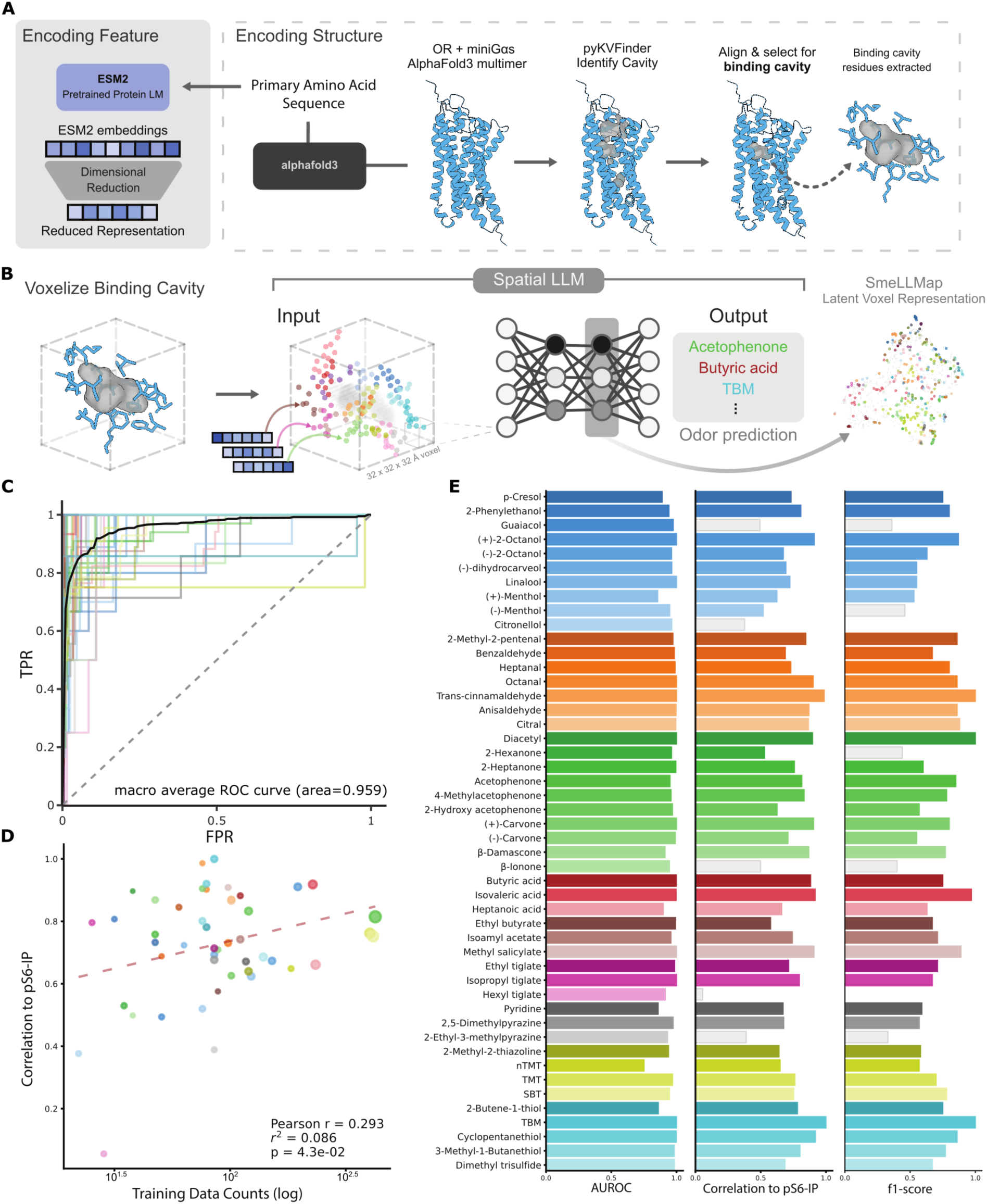
Spatial LLM architecture and model performance. **A.** Schematic of the data pre-processing pipeline. For sequence encoding, full-length OR sequences were input to ESM-2 to generate residue-wise embeddings, which were subsequently reduced in dimensionality by principal component analysis (PCA). In parallel, the same sequences were used to predict OR structures (OR+miniGαs) using AlphaFold3. Predicted structures were analyzed with pyKVFinder to identify internal cavities, which were filtered against predefined canonical binding cavity coordinates. Residues surrounding the retained binding cavity were extracted to generate a structural representation for each OR. **B.** Spatial LLM modeling pipeline. OR structural representations were voxelized into three-dimensional grids. Voxel occupancy encoded spatial structure, while reduced ESM-2 features were assigned to voxels based on residue identity, yielding a spatially and feature-aware representation of each OR. These voxel tensors were used as input to a convolutional neural network (CNN) trained to predict binary OR activation labels derived from pS6-IP activation z-scores. The output of the final fully connected layer was additionally extracted as a latent embedding, termed ‘smeLLMap’, representing learned receptor features. **C.** Receiver operating characteristic (ROC) curve summarizing CNN classification performance across all OR–odorant pairs with a macro-averaged curve plotted in black. **D.** CNN model prediction correlation to pS6-IP labels for individual odorants, plotted against the number of OR-ligand pairs in training data. Circle size is proportion to the number of test-set molecules. **E.** Model performance metrics evaluated per odorant, including area under the ROC curve (AUROC), correlation between prediction and pS6-IP activation values, and F1 score.

The resulting cavity-centered receptor structures were discretized into a three-dimensional voxel grid of size 32 x 32 x 32 at 1 Å resolution, with the grid centered on the predicted ligand-binding cavity (Fig. 3B). For each receptor, residues contributing to the binding cavity were first identified based on their spatial proximity, and each of their amino acid residue’s atomic coordinates were mapped onto the voxel grid according to their three-dimensional positions. Each occupied voxel was annotated with sequence-derived features obtained from ESM2 embeddings corresponding to the residue mapped to that spatial location. This voxelization strategy yields a spatially aligned, feature-aware representation of each receptor that jointly encodes binding cavity structure and residue identity for three-dimensional convolutional learning.

To provide functional supervision for training, we leveraged the pS6-IP-Seq data described above. Using the activation z-score metric greater than or equal to 2, which equates to roughly 0.05 of population in normal distribution, each OR was assigned to binary activated or inactive for each individually tested odors. This labeling scheme transforms the experimental response data into a multi-label classification task: given the structural and embedding representation of an OR, predict which odor it is likely to respond to. In this way, the pipeline effectively provides the first attempt at building an interpretable olfactory map directly from structural information.

### Convolutional Neural Network Odor Response Model

To predict OR odor responses, we trained a convolutional neural network (CNN) using voxelized binding cavity representations of 349 ORs responding to at least one odorant. For each receptor, five independent AlphaFold3 structural predictions were included, and voxel-level data augmentation was applied to increase robustness to minor structural variation. The dataset was randomly partitioned into training, validation, and test sets using a 70/15/15 split at the receptor level to prevent information leakage across folds. The model was optimized using the Adam optimizer with binary cross-entropy loss in a multi-label classification setting, reflecting the fact that individual receptors may respond to multiple odorants.

Across training runs, both training and validation losses decreased with no evidence of divergence between the two curves, indicating stable optimization without overfitting (fig. S4). Improvements in loss were accompanied by concordant increases in validation F1-score, demonstrating that optimization gains translated into improved classification performance rather than trivial fitting to dominant classes (fig. S4).

The final Spatial LLM CNN achieved consistently high predictive performance across the odor panel. Multi-class ROC analysis yielded a macro-averaged AUCROC of 0.959, indicating strong overall discriminative ability across odors (Fig. 3C). However, because the dataset is heavily class-imbalanced, with substantially more inactive than active OR–odorant pairs, ROC–AUC alone can overestimate performance due to the abundance of true negatives (*40*). To address this potential limitation, we evaluated additional performance metrics that are more sensitive to prediction confidence and class balance. First, we assessed whether model output probabilities reflected experimentally measured activation strength by correlating predicted confidence scores with pS6-IPseq activation values across receptors for each odorant (Fig. 3D). This analysis revealed strong correspondence between model confidence and biological activation, with 41 of 48 odorants exhibiting correlation coefficients greater than 0.6 and a mean correlation of 0.71 across odors (Fig. 3E and table S1). Second, precision–recall performance remained robust, with 32 of 48 odorants achieving F1-scores above 0.6 and a mean F1-score of 0.67, indicating balanced sensitivity and precision across most odor categories (Fig. 3D, fig. S4 and table S1).

We further examined the relationship between predictive performance and the number of positive training ORs available per odorant. Correlation strength increased with the number of activated ORs used for training, revealing a significant positive association between data availability and model performance (Fig. 3D). Notably, this trend was not deterministic, several odorants with relatively few positive examples nonetheless achieved high correlation and F1-scores, suggesting that certain odor response patterns are more easily inferred or generalized between receptors. Together, these complementary assessments confirm that the high ROC–AUC values reflect discriminative performance rather than artifacts of class imbalance. The Spatial LLM framework therefore provides an approach for predicting OR–odorant interactions from structural representations.

### Feature Embeddings

To interpret the learned representation of OR–odorant interactions, we extracted receptor embeddings from the final fully connected layer of the trained CNN. These vectors provide compact, high-level encodings of receptor structure after transformation through the convolutional architecture. Embeddings for each OR were retained alongside experimentally determined odor response labels for downstream analyses.

We first examined the global organization of the learned representation using UMAP dimensionality reduction for visualization. The resulting two-dimensional embedding, which we term smeLLMap, provides a spatial map of the OR landscape in which each point corresponds to a single receptor and its position reflects similarity in the learned odor–response representation, with colors indicating ligand selectivity (Fig. 4A and fig S5). At a global level, receptors do not strongly segregate by canonical family assignments (Fig. 4B). Consistent with this, functional organization emerges as the dominant structure. When receptors responding to individual odorants are visualized, they form distinct and coherent clusters within smeLLMap (Fig. 4C and fig. S5), indicating that the learned representation groups ORs according to shared ligand selectivity rather than phylogenetic relatedness. To quantify the relationship between embedding proximity and sequence similarity, we computed Grantham distance between 5 k nearest neighbors (kNN) derived from smeLLMap in comparison with those defined by amino acid-based neighbors for each ORs. Nearest neighbors in the smeLLMap exhibited significantly higher Grantham distances than amino acid derived neighbors (Fig. 4B). This result demonstrates that local organization in the smeLLMap is not solely driven by sequence similarity, but instead reflects alternative features captured by the model.

**Figure 4.**
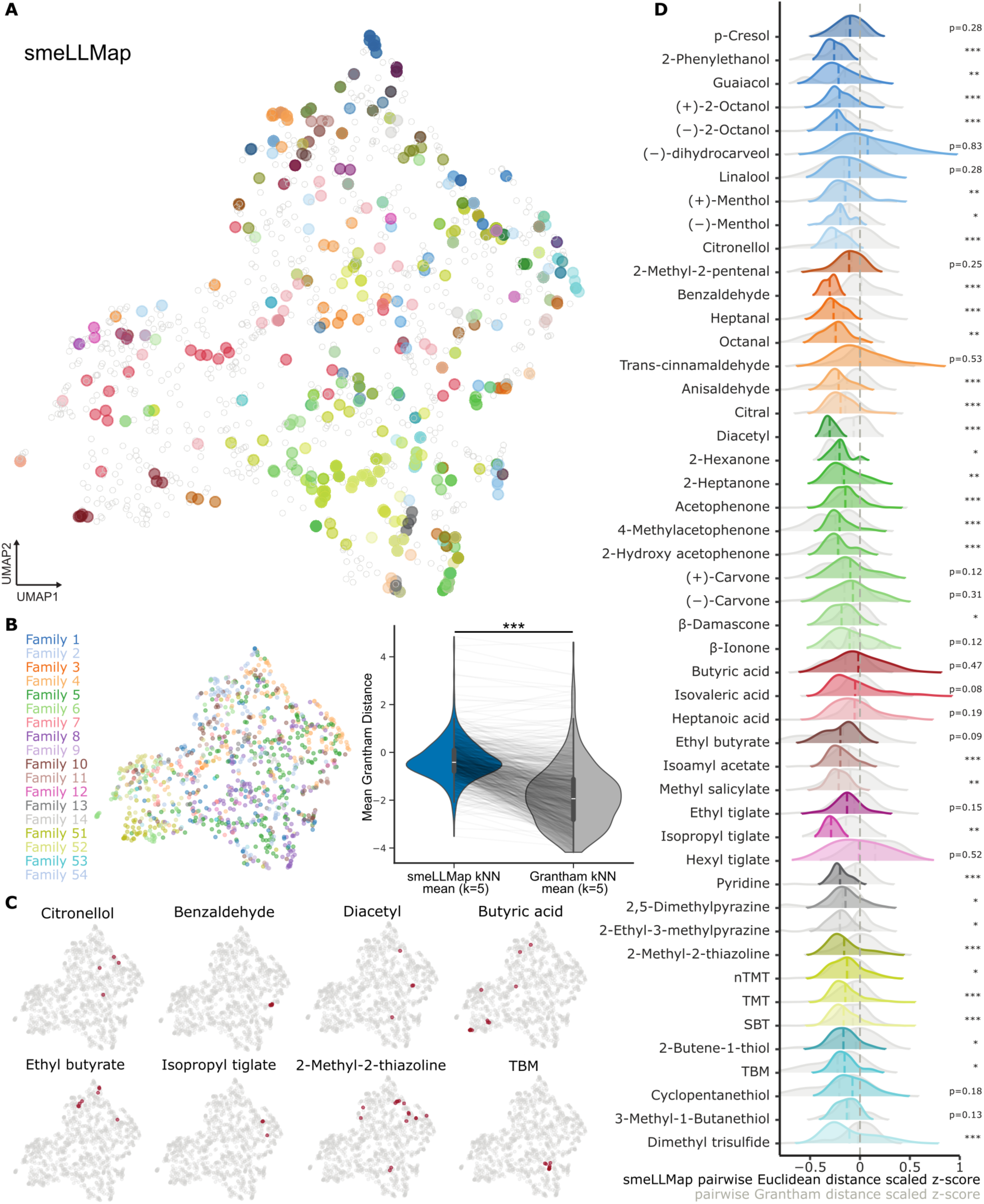
SmeLLMap captures odor-specific relationships among odorant receptors. **A.** Left: smeLLMap visualization of OR embeddings using UMAP dimensionality reduction. Each point represents a single OR; receptors with no detected responses are shown in hollow grey, while responding receptors are overlayed by odorant colors to which they respond based on pS6-IP activation labels. Right: individual smeLLMap subplots highlighting ORs response to a single odorant. **B.** Left: smeLLMap representation of ORs by family. Right: Violin plots show Grantham distance between neighbor distributions across receptors in smeLLMap. line indicates matched random pairs. Paired t-test, *p* = 1.05 × 10⁻^35^. **C.** smeLLMap representation of individual ORs activated (labeled in red) by specific odorant clusters. **D.** Violin plots of pairwise Euclidean distances between CNN-derived embeddings of ORs responding to the same odorant. Statistical significance was assessed using permutation tests with randomly sampled OR groups matched for group size per odorants. Grey violins denote the corresponding pairwise Grantham distance distributions for the same OR sets.

We next asked whether model embedding-based similarity better reflects shared ligand selectivity than primary amino acid sequence similarity. To address this, we computed pairwise Grantham distances between ORs responding to the same odor and compared these distributions with Euclidean distances measured in the smeLLMap. Subsequently, for 41 of 48 odorants, responding ORs show higher similarity between learned embedding than amino acid similarity (Fig. 4D and table S2). Together, these patterns and difference indicates that the learned embedding representation captures functional similarities between receptors that are not apparent from primary amino acid sequence similarity alone, highlighting the added value of the CNN-derived structural embedding over sequence-based metrics.

### Chemical and Perception Inference

We next asked whether the smeLLMap reflects relationships between odorants, despite the model having no explicit access to ligand chemical features during training. To address this, we first examined whether receptors responding to chemically similar odorants occupy similar regions in the learned latent space. For each odorant, the set of responding OR embeddings was treated as a distribution, and pairwise distances between odorants were quantified using Wasserstein distance (Fig. 5A and fig. S6). Odorants with high structural similarity exhibited significantly smaller distances between their corresponding receptor distributions. This was observed for closely related compounds such as stereoisomeric pairs, as well as homologous series including heptanal and octanal, which differ by a single carbon unit (Fig. 5A and fig. S6). In contrast, chemically dissimilar odorants, such as octanal and (+)-carvone, showed substantially larger distances between their respective receptor distributions. These results indicate that smeLLMap can infer chemical similarity through receptor organization, even in the absence of explicit ligand descriptors.

**Figure 5.**
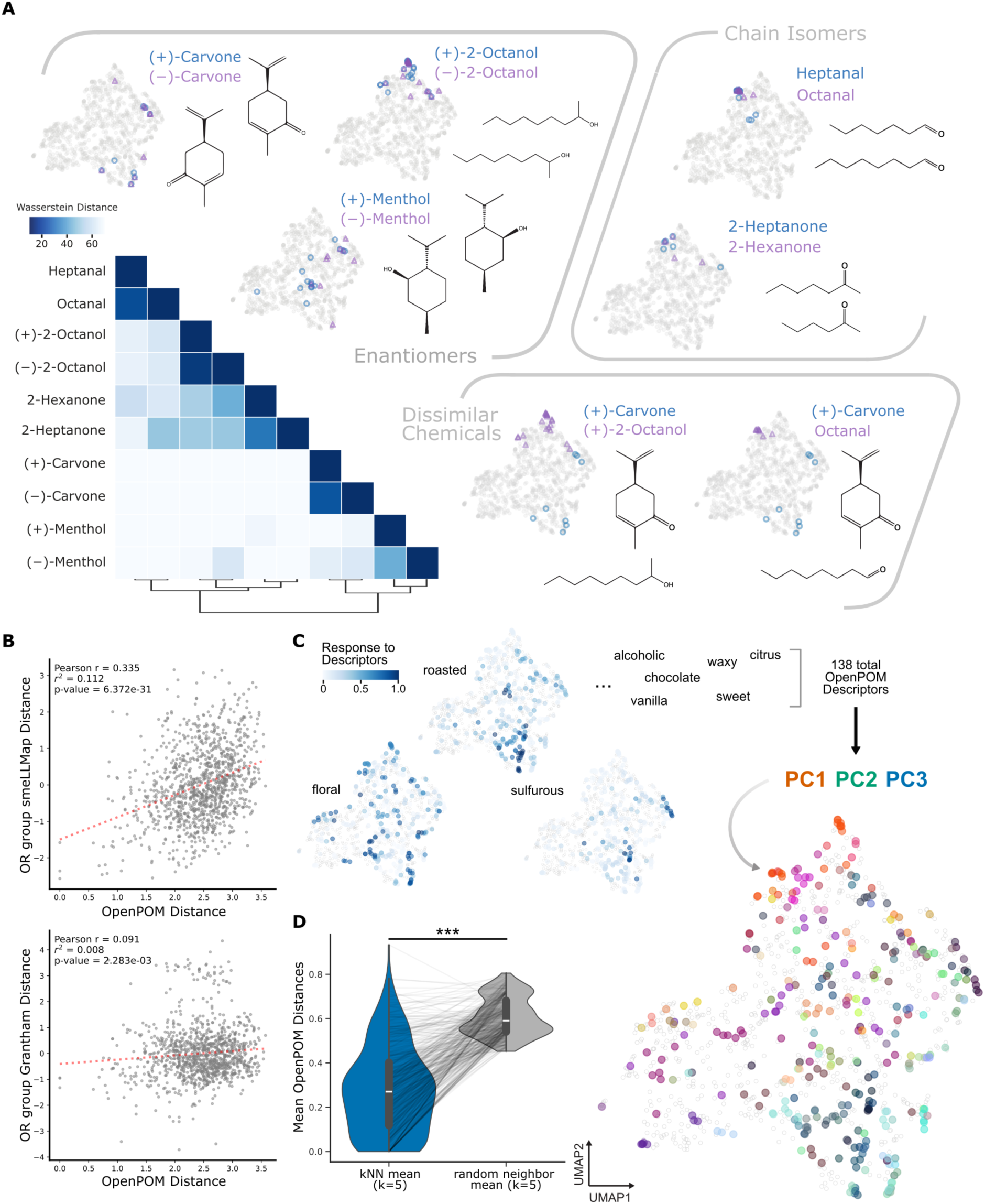
SmeLLMap captures chemical and perceptual similarity relationships among odorant receptors. **A.** smeLLMap reflects chemical relationships between odorants. Left: Heatmap of Wasserstein distances between distributions of OR embeddings for each odorant, clustered by similarity. Right: Representative smeLLMap projections illustrating receptor distributions for selected odorant pairs, including enantiomers, structurally related compounds (heptanal vs. octanal; 2-heptanone vs. 2-hexanone), and chemically dissimilar odorants ((+)-carvone vs. octanal). **B.** Quantitative comparison between receptor embedding derived distances and perception-informed chemical space. Top: Pearson correlation between pairwise Wasserstein distances of responding OR groups in smeLLMap and corresponding Euclidean distances in OpenPOM space across all 48 odorants. Bottom: Equivalent analysis using Grantham sequence distances in place of embedding distances, showing reduced correspondence with OpenPOM-derived relationships. **C.** Mapping perceptual features onto receptor embedding space. Left: Selected OpenPOM descriptors (e.g., floral, roasted, sulfurous) projected onto smeLLMap, revealing spatially localized receptor tuning. Right: Integration of all 138 OpenPOM descriptors, reduced by principal component analysis and visualized as RGB color channels (PC1–PC3). **D.** Local neighborhood structure in embedding space reflects perceptual similarity. For each receptor, the mean OpenPOM Euclidean distance of its five nearest neighbors in smeLLMap is compared to a random expectation. Violin plots show distributions across receptors; line indicates matched random pairs. Paired t-test, *p* = 6.46 × 10⁻^96^.

While these relationships highlight the ability of the model to recover chemical similarities between enantiomer and chain isomer odorants, we next asked whether this organization extends beyond chemical features to reflect perceptual similarity. To test this, for each pair of odorants we computed the distance between their responding OR distributions in smeLLMap and compared these values to the corresponding distances derived from OpenPOM (*21, 34*) space. Across all 48 odorants, we observed a significant positive correlation between receptor embedding distance and perceptual distance (*r*^2^=0.112, *p*=6.372e-31) (Fig. 5B and fig. S6A), indicating that odorants perceived as more similar by humans panels tend to be represented by similar ORs represented with regions of receptor space. Notably, this relationship was not clear when using amino acid sequence-based Grantham distances between responding ORs (*r*^2^=0.007, *p*=2.283e-3) (Fig. 5B and fig. S6B), demonstrating that primary sequence similarity alone does not capture perceptual organization.

To further visualize how perceptual features are represented across receptor space, we projected individual OpenPOM-derived odor descriptors onto smeLLMap. By mapping specific odor descriptors, such as floral, roasted and sulfurous qualities, revealed localized clusters of receptors preferentially associated with descriptors (Fig. 5C). We next examined whether this organization could be observed at a global level by integrating the full set of 138 OpenPOM descriptors. Utilizing dimensionality reduction of these perceptual features followed by projection onto smeLLMap, with each principal component axis as a representation RGB color space revealed further clustering of colored gradients. Likely indicating that perceptual features may also be organized and inferred by the OR space, highlights a structured encoding in which different regions of receptor space are tuned to distinct combinations of perceptual features. To quantitatively assess whether local neighborhoods in smeLLMap reflect shared perceptual tuning, we computed kNN relationships and compared the OpenPOM (*21, 34*) representations of neighboring receptors. Because our analyses relate mouse receptor activation to human perception derived features, they should be interpreted with appropriate caution. Nevertheless, receptors exhibited significantly greater similarity in perceptual representation to their nearest neighbors than random (Fig. 5D), supporting the idea that smeLLMap captures perceptually relevant organization.

Together, these results demonstrate that smeLLMap captures not only chemical similarity but also perceptual relationships between odorants. This organization reflects encoding in which receptor structure gives rise to functional groupings that align with both molecular features and human odor perception. This finding suggests that perceptual similarity may emerge, at least in part, from the structure of receptor activation patterns encoded within the OR binding cavity.

### Feature Importance

While the smeLLMap captures the global organization of receptor–odorant relationships, we next sought to identify which specific structural features most strongly influenced individual model predictions. To this end, we applied Integrated Gradients (IG), which attributes a model’s output to input voxels by measuring how the prediction changes as structural information is gradually introduced from a baseline representing the absence of receptor structure. Voxels with large attribution magnitudes, either positive or negative, exert the strongest influence on the prediction, yielding a three-dimensional importance map over the binding cavity that highlights features that promote or disfavor ligand binding (Fig. 6A). Importantly, each voxel in our representation encodes both spatial location and residue-level biochemical features. As a result, voxel attribution reflects the joint contribution of three-dimensional positioning and physicochemical identity, rather than isolating either factor in abstraction. This integrated representation is advantageous, as ligand recognition in ORs depends on the precise spatial arrangement of chemically specific residues within the binding pocket. Attribution scores therefore capture how geometry and chemistry act together to shape receptor selectivity (Fig. 6A).

**Figure 6.**
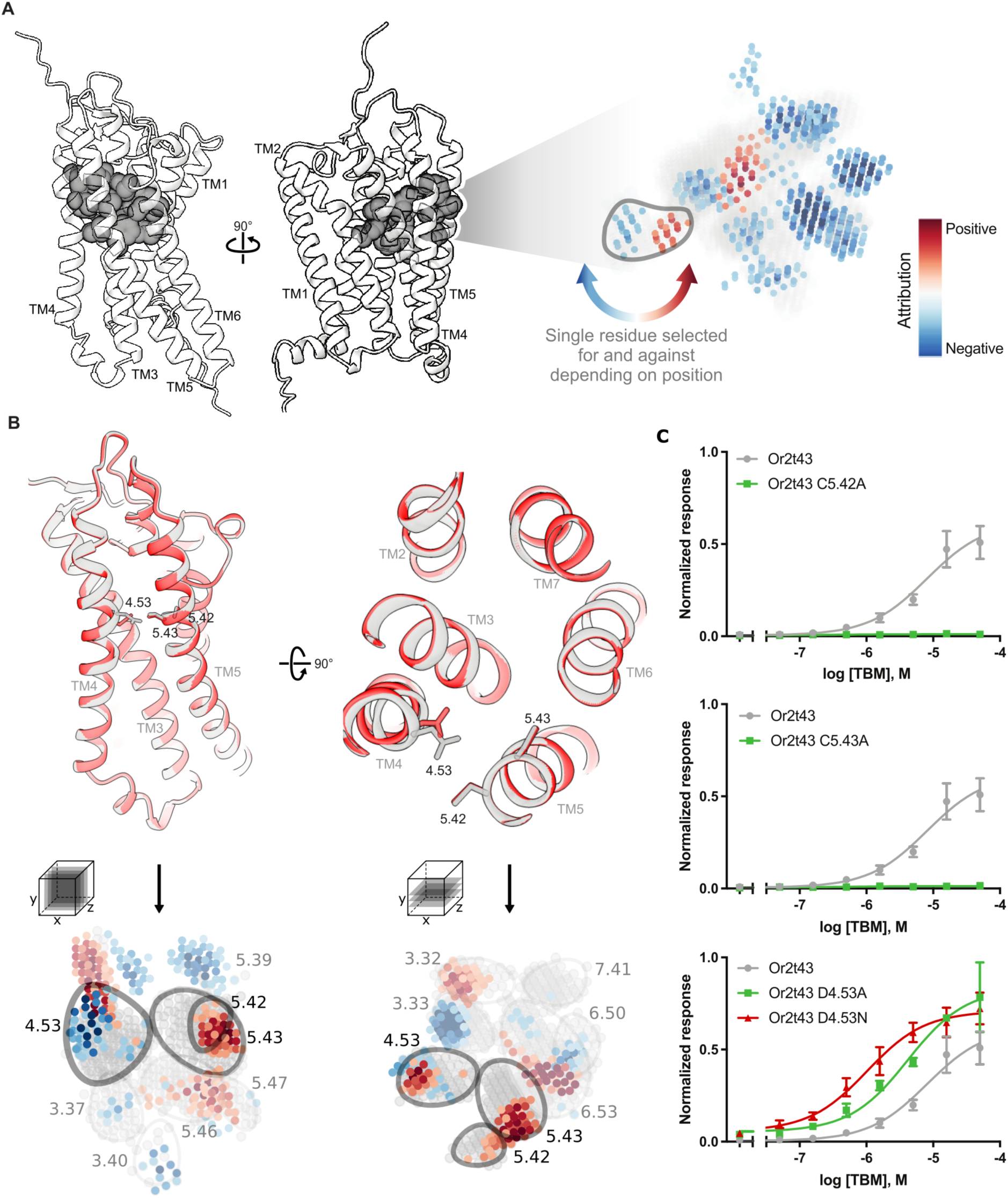
Voxel-level attribution identifies structural determinants of odor selectivity. **A.** Structural and voxel-based representations of olfactory receptor binding cavities. Left, occupancy representation of amino acid residues defining the extracted canonical binding cavity mapped onto the OR structure, shown from two viewing angles. Right, corresponding three-dimensional voxel saliency map, highlighting spatial regions within the cavity that contribute most strongly to model predictions. **B.** Top: Predicted AlphaFold3 structures of native Or2t43 and Or2t43D4.43N mutant with side and top view orientation. Bottom: Stacks aggregated saliency maps for receptors responding to tert-butyl mercaptan (TBM). Left and middle, orthogonal slice views through the averaged voxel attribution map reveal conserved high-importance regions within the binding cavity. **C**. functional validation by mutagenesis of residues D4.53, C5.42, and C5.43 in Or2t43, assessed using a real-time cAMP luciferase assay. Data are shown as mean ± s.d. from *n* = 3 independent experiments.

Notably, when attributions were aggregated across multiple receptors responding to the same odorant, importance did not simply track residue identity. Instead, the structural conformation of those residues appeared to influence their contribution to ligand binding. The same residue position could contribute positively or negatively depending on whether its side chain projected inward to constrict the cavity or outward to expand it, likely excluding or permitting ligand access (Fig. 6A). Together, these results indicate that odorant selectivity is governed not only by which residues are present, but by how their three-dimensional positioning modulates cavity accessibility.

We next asked whether this combined embedding and voxel-level attribution framework could be used to guide odorant-receptor activation. As a case study, we focused on tert-butyl mercaptan (TBM), a sulfur-containing odorant widely used as a natural gas additive due to its strong detectability.

Guided by the IG attributions, we engineered targeted mutations in Or2t43 and assessed their responses to TBM (Fig. 6B). Although Or2t43 is robustly activated by TBM in its native form, mutation of either C5.42 or C5.43 resulted in a complete loss of activation, consistent with disruption of the predicted sulfur-binding coordination site. In contrast, substitution of D4.53 with alanine, reducing side-chain size and polarity, or with asparagine, maintaining steric occupancy while altering charge, both increased TBM responsiveness to TBM. These reciprocal effects are consistent with the directionality of the model’s attribution scores and indicate that the highlighted residues exert functionally meaningful influence on ligand sensitivity. In addition, the model identified residue D4.53 as negatively associated with TBM responsiveness, suggesting a potential inhibitory role at this position.

To further examine the structural basis of these effects, we generated AlphaFold3 models of the corresponding Or2t43 mutants. Comparison of the predicted structures revealed that mutation of D4.53 to asparagine results in a repositioning of the side chain within the binding cavity. Notably, the mutated N4.53 side chain extends into a spatial region immediately adjacent to, but distinct from, the original asparagine position, consistent with the observed increase in TBM responsiveness. These structural changes align with the attribution-derived importance map, providing support that local rearrangements within the binding cavity can modulate ligand sensitivity in a predictable manner.

Together, these results demonstrate that our interpretable Spatial LLM framework not only captures the global organization of odor–receptor relationships but also provides mechanistic insight at residue-level resolution. By linking structural context, learned embeddings, and voxel-level attribution, the model enables rational prediction of functionally relevant mutations and supports targeted modulation of receptor sensitivity to specific odorants.

## Discussion

This study demonstrates that OR–ligand selectivity can be learned and predicted through a structure-centered framework that explicitly models the three-dimensional organization and sequence-informed composition of the receptor binding cavity. By integrating in vivo pS6-IP-Seq activation measurements with spatial learning, we show that the canonical binding cavity contains sufficient information to organize receptor responses across chemically diverse odorants (Fig. 3B,D and 4D). These findings indicate that the determinants of ligand selectivity are encoded within the binding pocket itself, even without explicit modeling of alternative or allosteric interaction sites, thereby addressing a long-standing challenge in linking receptor variation to interpretable chemical recognition.

A key conceptual advance of this work is the demonstration that functional similarity among ORs is better captured by learned structural embeddings than by primary sequence similarity alone (Fig. 4B). While traditional sequence-based metrics and protein language model embeddings reflect evolutionary relationships, they fail to fully explain why distantly related receptors can converge on similar ligand selectivity. In contrast, the CNN-derived embedding organizes receptors according to shared response profiles, revealing functional neighborhoods that spans beyond phylogenetic relationship. This result suggests that ligand selectivity is governed by higher-order structural constraints, such as cavity geometry and residue positioning, that are not apparent from sequence comparisons.

Notably, this organization extends beyond chemical similarity to reflect perceptual relationships between odorants (Fig. 5A-D and fig. S6). Despite being trained without explicit ligand features, the learned embedding correlates with perceptual distances derived from OpenPOM (*34*), a model trained on chemical molecules to predict odor perception, whereas amino acid sequence-based distances do not (Fig. 5B). This correspondence suggests that receptor binding cavity encode features that can be used to infer odor perceptual similarity. At the same time, these comparisons should be interpreted with appropriate caution. The receptor structures and activation data in this study are derived from mouse, whereas OpenPOM reflects human perceptual judgments. Rather than implying a direct mapping between mouse receptor activity and human perception, this result points to shared, cross-species constraints in how chemical features are transformed into receptor-level representations. Such constraints may arise from conserved structural principles governing receptor–ligand interactions. Future work integrating human OR repertoires, alongside matched perceptual datasets, will be important to determine the extent to which receptor-level features directly encode perceptual dimensions. Establishing this link will help clarify whether the observed alignment reflects a generalizable organizing principle or emerges from partial overlap between species-specific receptor spaces.

Importantly, the interpretability of this representation allows functional predictions to be traced back to specific structural features (Fig. 6A-C). Integrated Gradients attribution revealed that both positively and negatively contributing voxels shape ligand responsiveness, highlighting residues that promote binding as well as those that disfavor activation. The observation that identical residues can exert opposite effects depending on their three-dimensional orientation underscores that odorant recognition is fundamentally a spatial phenomenon (Fig. 6A). These results support a model in which receptor tuning arises from the precise arrangement of residues within the binding cavity, rather than from residue identity alone.

Given the dynamic nature of OR activation, the ability of static AlphaFold3 predicted structures to support prediction and attribution is notable. While receptor conformational and ligand-induced rearrangements undoubtedly contribute to signaling, our results indicate that determinants of selectivity are already encoded in the active-state architecture of the binding pocket structure and ESM features (Fig. 3E). Although this framework does not explicitly model OR secondary or allosteric binding sites, the predictive performance suggests that such interactions, when present, either act through modulation of the canonical cavity or play a secondary role in determining primary ligand selectivity. Alternatively, the current framework may also suggest the frequency of binding outside of canonical binding cavity occur in relatively limited subset of receptors such that the overall statistical framework remains significant.

Additionally, by encoding amino acid features with ESM embeddings may also enables the feature to capture long-range dependencies that could be outside of the selected binding cavity itself (fig. S2). Future incorporation of receptor dynamics, alternative conformational states, or ligand-induced structural shifts may further refine predictions, but may not be required to recover core selectivity principles.

Beyond mechanistic insight, this study provides a scalable strategy for interrogating OR function at a systems level. The use of pS6-IP-Seq, coupled with an activation metric designed to minimize transcript abundance and variance biases, enabled broad and relatively unbiased sampling of receptor–odorant interactions in vivo (Fig. 1A-F). The resulting dataset supports a model that generalizes across receptors and odorants, including cases with sparse activation data (Fig. 3D), highlighting the importance of carefully designed functional metrics in enabling downstream computational modeling.

While the present study focuses on single odorants at fixed concentrations, several extensions could substantially broaden its scope. Expanding chemical coverage would improve representation of sparsely sampled ligand classes, while incorporating concentration dependence could reveal how cavity features govern sensitivity and dynamic range. Natural olfactory stimuli are typically complex mixtures, and extending the framework to model mixture responses or competitive interactions between ligands represents an important next step. Although caution is warranted when interpreting pS6-IP-Seq data, as odorants used for stimulation may be metabolized within the olfactory mucus before binding to ORs, integrating explicit ligand representations alongside receptor structures may further enable dissection of receptor–ligand complementarity and cross-reactivity.

The implications of this work extend beyond olfaction. ORs belong to the GPCR superfamily, which encompasses a wide range of receptors involved in sensory perception, neurotransmission, and pharmacological signaling. Demonstrating that structural features alone can organize receptor function in a highly diverse GPCR family suggests that similar approaches may be applicable to other receptors lacking extensive experimental characterization. In this context, the framework introduced here provides a generalizable strategy for linking receptor structure to ligand selectivity in an interpretable and predictive manner.

In summary, the central of this study is that OR function is encoded in the three-dimensional structure of the binding cavity in a way that can be learned, interpreted, and experimentally validated. By incorporating sequence-centric descriptions to structure-informed representations, this work contributes toward a predictive, mechanistic understanding of chemical sensing. As structural models, functional data, and computational methods continue to converge, this approach lays the groundwork for quantitative and ultimately generative models of olfactory coding.

## Data, code, and materials availability

All code used for data processing, structural modeling, and machine learning is publicly available at GitHub: https://github.com/Justice-Lu/OR_learning/tree/master. Raw and processed RNA-sequencing data generated in this study have been deposited in the NCBI Gene Expression Omnibus (GEO) under accession number GSE185415. Additional data supporting the findings of this study are available from the corresponding author upon reasonable request.

## Supplementary Materials

Materials and Methods

Supplementary Text

Figs. S1 to S6 Tables S1 to S2

