## Supplemental materials for "Decoding Smell from Receptor Structure"

**The PDF file includes:**

Materials and Methods
Figs. S1 to S6
Tables S1 to S2

#### Materials and Methods

##### Phosphorylated S6 ribosomal immunoprecipitation sequencing (pS6-IP-Seq)

Mice used for pS6-IP were ~3 weeks old, mixed sex, and littermates. Mice were killed by $CO_2$ asphyxiation and cervical dislocation. Olfactory tissue was rapidly dissected in Buffer B (2.5 mM HEPES KOH pH 7.4, 0.63% glucose, 100 μg/mL cycloheximide, 5 mM sodium fluoride, 1 mM sodium orthovanadate, 1 mM sodium pyrophosphate, 1 mM β-glycerophosphate, in Hank's balanced salt solution). Tissue pieces were then minced in 1.35 mL Buffer C (150 mM KCl, 5 mM $MgCl_2$, 10 mM HEPES KOH pH 7.4, 0.100 μM Calyculin A, 2 mM DTT, 100 U/mL RNAsin, 100 μg/mL cycloheximide, protease inhibitor cocktail, 5 mM sodium fluoride, 1 mM sodium orthovanadate, 1 mM sodium pyrophosphate, 1 mM β-glycerophosphate)and subsequently transferred to homogenization tubes for steady homogenization at 250 rpm three times and at 750 rpm nine times at 4 °C. Samples were then transferred to a 1.5 mL LoBind tube (Eppendorf 022431021) and clarified at 2000xg for 10 min at 4 °C. The low-speed supernatant was transferred to a new tube on ice, and 90 μL of NP40 (Sigma 11332473001) and 90 μL of 1,2-diheptanoyl-sn-glycero-3-phosphocholine (DHPC, Avanti Polar Lipids 850306P, 100 mg/0.69 mL) were added to this solution. This solution was mixed and then clarified at a max speed (17,000xg) for 10 min at 4 °C. The resulting high-speed supernatant was transferred to a new tube where 20 μL was saved and transferred to a tube containing 350 μL buffer RLT. To the remainder of the sample, 1.3 μL of 100 μg/mL cycloheximide, 27 μL of phosphatase inhibitor cocktail (250 mM sodium fluoride, 50 mM sodium orthovanadate, 50 mM sodium pyrophosphate, 50 mM β-glycerophosphate) and 6 μL of anti-pS6 antibody (Cell Signaling

D68F8) were added. The sample was gently rotated for 90 min at 4 °C. To prepare beads, 100 μL of beads (Invitrogen 10002D) was washed three times with 900 μL of buffer A (150 mM KCl, 5 mM $MgCl_2$, 10 mM HEPES KOH pH 7.4, 10% NP40, 10% BSA), and once with 500 μL of buffer C. Sample homogenate was added to the beads and incubated with gentle rotation for 60 min at 4 °C. Following incubation, beads were washed with four times with 700 μL of buffer D (350 mM KCl, 5 mM $MgCl_2$, 10 mM HEPES KOH pH 7.4, 10% NP40, 2 mM DTT, 100 U/mL RNAsin, 100 μg/mL cycloheximide, 5 mM sodium fluoride, 1 mM sodium orthovanadate, 1 mM sodium pyrophosphate, 1 mM β-glycerophosphate). During the final wash, beads were moved to room temperature, wash buffer was removed, and 350 mL of buffer RLT was added. Beads were incubated in buffer RLT for 5 min at room temperature. Buffer RLT containing immunoprecipitated RNA was then eluted and stored at −80 °C until clean up using a kit (Qiagen 74004). cDNA was generated using 11 rounds of amplification with 10 ng RNA input. DNA libraries were prepared using a half-sized Nexterra XT DNA Library Preparation Kit (Illumina 15032354) protocol as per the manufacturer's guidelines. Libraries were sequenced on either HiSeq 2000/2500 (50 base pair single read mode) or NextSeq 500 (75 base pair single read mode) with 6–12 pooled indexed libraries per lane.

**RNA-Seq alignment, quantification, and differential expression analysis**

Reads were aligned against a modified GRCm38.p6 (M25) reference, in which we deleted ENSMUSG00000116179 (Olfr290), using STAR (*1*) with --outFilterMultimapNmax 10. Reads mapping to Olfr290 were inferred from ENSMUSG00000070459, with the rationale that this gene model included ENSMUSG00000116179 plus untranslated regions. Gene-level read quantification was done using RSEM (*2*). Differential expression analysis was performed against all genes using EdgeR (*3*). Gene nomenclature was retrieved from BioMart (*4*). Intact Olfr genes with identifiable sequences were filtered, and p-values were then re-corrected by FDR. Only ORs exhibiting odor response to at least one of the tested odorants ($\log_2 FC > 0$ and $FDR < 0.05$) were considered. A total of 555 ORs responded across the 72 different odorants at various concentrations. A total of 375 ORs were responsive to unique odorants at the lowest tested concentrations. Raw and processed RNA-Seq datasets generated as part of this study are available from NCBI GEO at accession GSE185415.

**Source of odorants**

The following odors and concentrations were used for molecular response profiling and model training: 1% p-Cresol (Sigma C85751), 1% 2-phenylethanol (Sigma 77861), 1% guaiacol (Sigma G10903), 10% (+)-2-octanol (Sigma O4504), 10% (−)-2-octanol (Sigma 147990), 1% linalool (Sigma L2602), 1 M (+)-menthol (Sigma 224464), 1 M (−)-menthol (Sigma M2780), 1% citronellol (Sigma W230915), 1% 2-methyl-2-pentenal (Sigma 294667), 1% benzaldehyde (Sigma W212717), 1% heptanal (Sigma W254002), 1% octanal (Sigma O5608), 1% trans-cinnamaldehyde (Sigma C80687), 0.01% anisaldehyde (Sigma A88107), 0.01% citral (Sigma W230316), 1% ethyl butyrate (Sigma W242713), 1% isoamyl acetate (Sigma 306967), 1% methyl salicylate (Sigma W274502), 1% diacetyl (Sigma W237027), 1% 2-hexanone (Sigma 103004), 1% 2-heptanone (Sigma 537683 (*5*)), 0.01% acetophenone (Sigma W200910), 1% 4-methylacetophenone (Sigma W267708), 1% (+)-carvone (Sigma 22070), 1% (−)-carvone (Sigma 22060), 1% β-damascone (Sigma W324300), 1% β-ionone (Sigma W259525), 1% pyridine

(Sigma 270970), 1% 2,5-dimethylpyrazine (Sigma 175420 (*5*)), 1% 2-ethyl-3-methylpyrazine (Sigma W315508), 0.01% 2-butene-1-thiol (CheMall Corp OR116574), 1% tert-butyl mercaptan (2-methyl-2-propanethiol; Sigma 109207), 0.01% cyclopentanethiol (Sigma W326208), 1% 3-methyl-1-butanethiol (Sigma W385808), 1% dimethyl trisulfide (Sigma W327506), 1% 2-methyl-2-thiazoline (Sigma M83406), 1% 2,4,5-trimethylthiazole (nTMT; Sigma 219185), 0.01% 2,4,5-trimethyl-4,5-dihydrothiazole (TMT, synthesized (*6*)), 0.01% 2-sec-butyl-4,5-dihydrothiazole (SBT; synthesized (*5*)), 1% ethyl tiglate (Sigma W246000), 1% isopropyl tiglate (Sigma W322903), and 1% hexyl tiglate (Sigma W500909).

**Chemical space estimation**

To estimate chemical space, we first identified 4680 small molecules commonly found in foods and fragrances from http://www.thegoodscentscompany.com/ (*7*). Smiles strings for these molecules and the 48 in the test odor set were then downloaded from PubChem, and Morgan-algorithm based fingerprints were calculated using Rdkit (*8, 9*). Chemical space was estimated by PCA dimensionality reduction on all molecules.

To complement this structure-based representation with a perceptual framework, we additionally embedded all molecules using OpenPOM(*10, 11*), a message-passing neural network trained to predict human odor perception from chemical structure. The resulting OpenPOM feature vectors provide a high-dimensional representation of perceptual odor space derived from human panel data (*12*). These embeddings were used to assess perceptual diversity of the odor panel, perform pairwise distance comparisons, and relate receptor embedding organization to perceptual similarity in downstream analyses.

**Receptor alignment and space estimation**

Mouse ORs were aligned to one another using the MAFFT E-INS-I method with manual refinements (*13*). The resulting alignment file was subjected to ModelTest-NG to identify ideal amino acid substitution models (*14*). Receptor pairwise similarity was calculated by summing amino acid differences at each position by Grantham's amino acid distances (*15*). Multidimensional scaling on all ORs.

**Structural Model Generation**

All mouse olfactory receptor (OR) protein sequences were compiled together with mini-Gα peptide definitions into standardized JSON configuration files. For each receptor, five independent structural predictions of the OR–miniGα complex were generated using AlphaFold-Multimer (*16*), capturing variability in receptor conformation under identical modeling conditions. In parallel, five ORs with respective ligands with available cryo-EM structures (OR51E2, consOR1, consOR2, consOR4, consOR51) were processed through the same workflow to serve as experimentally grounded structural references. Following structural prediction, all OR complexes were placed into a common coordinate frame using TM-align (*17*), ensuring consistent spatial orientation of transmembrane helices and intracellular domains across receptors. Internal solvent-accessible pockets were then identified using pyKVFinder (*18*), which outputs cavity coordinates. This procedure was applied uniformly to predicted models, cryo-EM structures, and later to ligand-bound complexes.

**Canonical Binding Cavity Identification**

To determine the biologically relevant ligand binding region, we incorporated receptor–odorant activation data derived from pS6-IP assays. For each experimentally activated receptor–ligand pair, AlphaFold3 was used to model receptor–miniGα–ligand complexes, followed by cavity detection with pyKVFinder. Ligand-occupied pockets were spatially compared across cryo-EM receptors, and regions demonstrating ≥50% overlap across ligand-bound cavities were defined as the canonical binding cavity. This region represents the reproducibly utilized ligand-engagement space across structurally diverse ORs.

The canonical binding cavity mask was then projected back onto ligand-free OR models for the entire repertoire. For each receptor, pyKVFinder-defined cavities were filtered relative to the canonical binding cavity; regions with <80% spatial overlap were discarded. This filtering process preserved a single, standardized binding cavity per receptor while removing allosteric, non-functional or receptor-specific peripheral pockets. Individual receptor's cavity surrounding amino acid then defined by less than 5 Å. The resulting structures consist of coordinates of each OR binding cavity and amino acid coordinates.

**Voxelization and Feature Encoding**

For each receptor, all residues spatially lining the canonical binding cavity were extracted to construct a localized structural representation. The cavity volume of each receptor was then discretized into a 32 × 32 × 32 voxel grid, maintaining 1Å spatial resolution, such that each voxel encodes a fixed physical region of three-dimensional space surrounding the binding pocket. For every voxel containing protein atoms, the residue occupying that spatial position was assigned as the voxel occupant. To provide a rich biochemical and evolutionary descriptor beyond atomic coordinates, the reduced ESM sequence embedding vector corresponding to that residue was inserted as the voxel feature representation. Voxels not occupied by residues were assigned a null background vector. This procedure resulted in a structured, spatially aligned, and biologically informed voxel map for each receptor. In this final representation, every OR is encoded as a voxelized binding cavity populated with residue-level embedding features, providing a standardized structural input for downstream convolutional neural network training and odor selectivity modeling.

**Heterologous luciferase assay**

Hana3A cells; which stably express $G_{olf}$, RTP1, RTP2, and REEP1; were grown in minimum essential medium eagle (MEM; Corning 10-010-CV) containing 10% Fetal Bovine Serum (FBS; vol/vol; Gibco 16000-044), penicillin-streptomycin (Sigma-Aldrich P4333), and amphotericin B (Gibco 15290018). Cells were cultured and incubated at 37°C, 5% $CO_2$, and saturated humidity for use with the Dual-Glo Luciferase Assay (Promega E2980)(*19, 20*). Cells were plated at 20-25% confluence on poly-D-lysine-coated 96-well plates (Corning 3843) overnight. After overnight incubation, cells were transfected with 6 mL of MEM containing 10% FBS, 0.5 µg SV40-RL (Promega E2980), 1 µg CRE-Luc (Promega E2980), 0.5 µg mouse RTP1s, 0.25 µg M3 muscarinic receptor, 0.5 µg of Rho-tagged receptor plasmid DNA, and 20 µg Lipofectamine 2000 (Invitrogen 11668019) per plate. Transfection medium was divided equally among the wells so that each OR-odorant combination could be conducted in triplicates. The following day, cells were incubated with 25µL of odorant solution diluted in CD-293 (Gibco

11913-019) containing 30 μM $CuCl_2$ (Sigma-Aldrich C-6641) and 2 mM glutamine (Gibco 25030-081) for 3.5 hours. cAMP-driven firefly Luciferase luminescence (Luc) was used to assess OR activation, and SV40-driven Renilla Luciferase luminescence (Ren) was used to control for variation in cell viability within wells. Cell luminescence was read by a POLARstar OPTIMA (BMG Labtech) luminometer, and normalized response values were calculated using the formula (Luc-400)/(Ren-400). ORs were considered responsive in vitro if ANOVA p-value was $< 0.05$ and ANOVA with post-hoc Dunnet's test correction p-adjusted was $< 0.05$ for at least 2 of the tested odor concentrations using the R package DescTools (v0.99.42). Log-logistic 4-parameter dose response curves were fit to the data using the R package drc (v3.0-1). In vitro responses were compared to in vivo responses by subtracting mean ligand-independent activity (luciferase values of ORs with no odor stimulation) from each of the ligand stimulated data points and summing. Scaled summed (+)-enantiomer responses were divided by scaled summed (-)-enantiomer responses and $\log_2$ transformed for comparison to $\log_2$FC (+)/(-) in vivo enrichments.

**Data analysis**

Analysis were performed in Python 3.8. Custom scripts for data analysis and visualization built using open-source python libraries (pandas, numpy, matplotlib, plotly, sklearn, scipy, seaborn and itertools). Scripts to replicate data analysis are available on github (https://github.com/Justice-Lu/OR_learning).

**Statistical analysis**

Experimental data are presented as mean ± standard deviation (SD), based on results from at least three independent experiments. Statistical analysis was conducted in Python 3.11.13 using SciPy v1.16.0. Paired comparisons (e.g., samples with different condition measured in the same experiment) were analyzed using the paired Student's t-test (ttest_rel). For unpaired comparisons between two independent groups, the non-parametric Wilcoxon rank-sum test (ranksums) was applied. A table with all the statistical comparison values and description can be found at Table 3. A two-tailed p-value less than 0.05 was considered statistically significant. Significance levels are denoted as follows: $p < 0.05$ (*), $p < 0.01$ (**), $p < 0.001$ (***), and $p \geq 0.05$ (n.s., not significant).

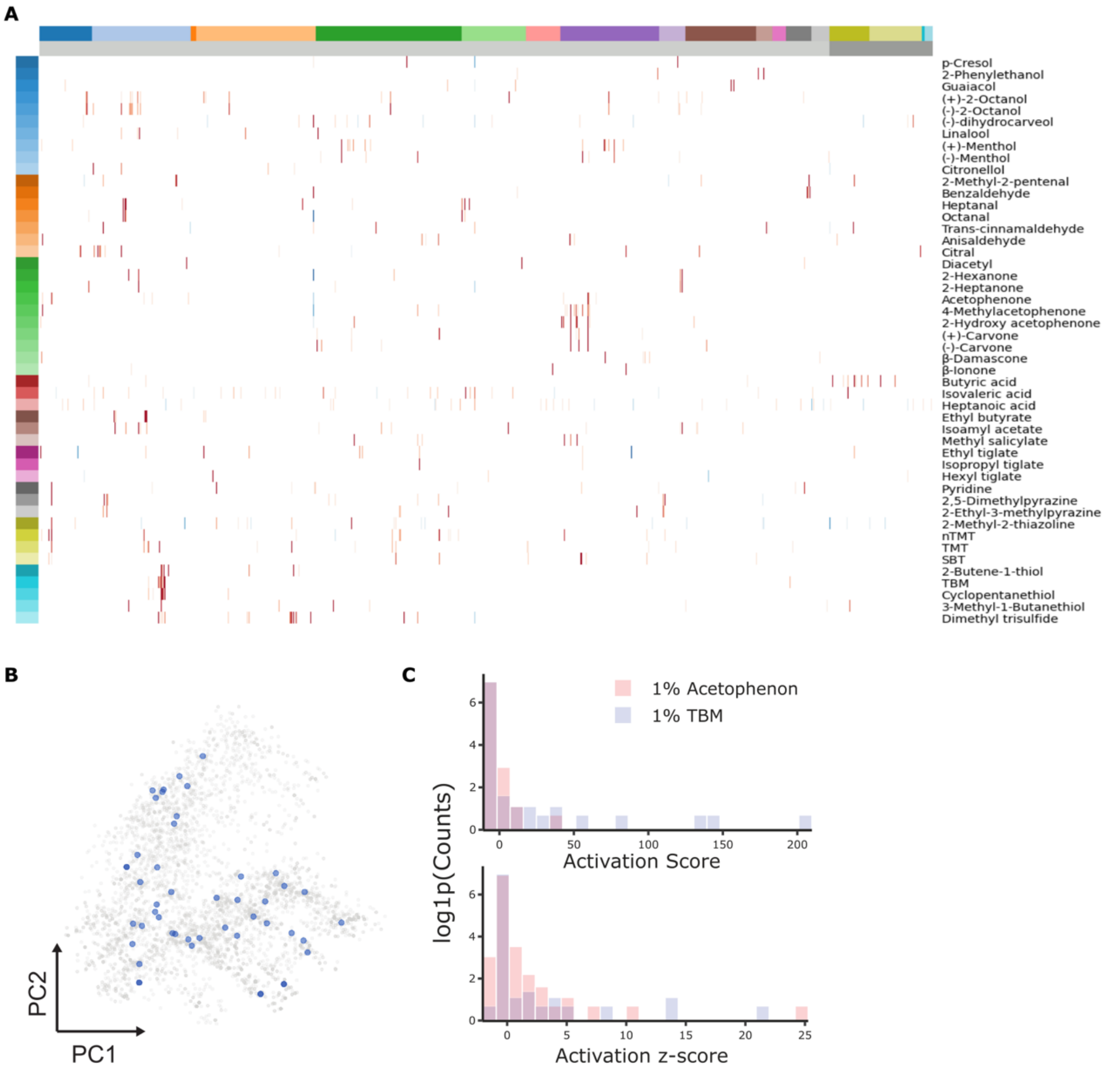


**Figure S1. Global overview of pS6-IP activation profiles and chemical space coverage.**

**A.** Heatmap of pS6-IP–derived activation z-scores across all tested odorant–receptor pairs. Rows correspond to the 48 odorants, and columns represent individual ORs, organized by class and family. **B.** Chemical space representation of odorants based on Morgan fingerprint principal component analysis (PCA). Odorants included in the pS6-IP panel are highlighted in blue relative to the full reference set.

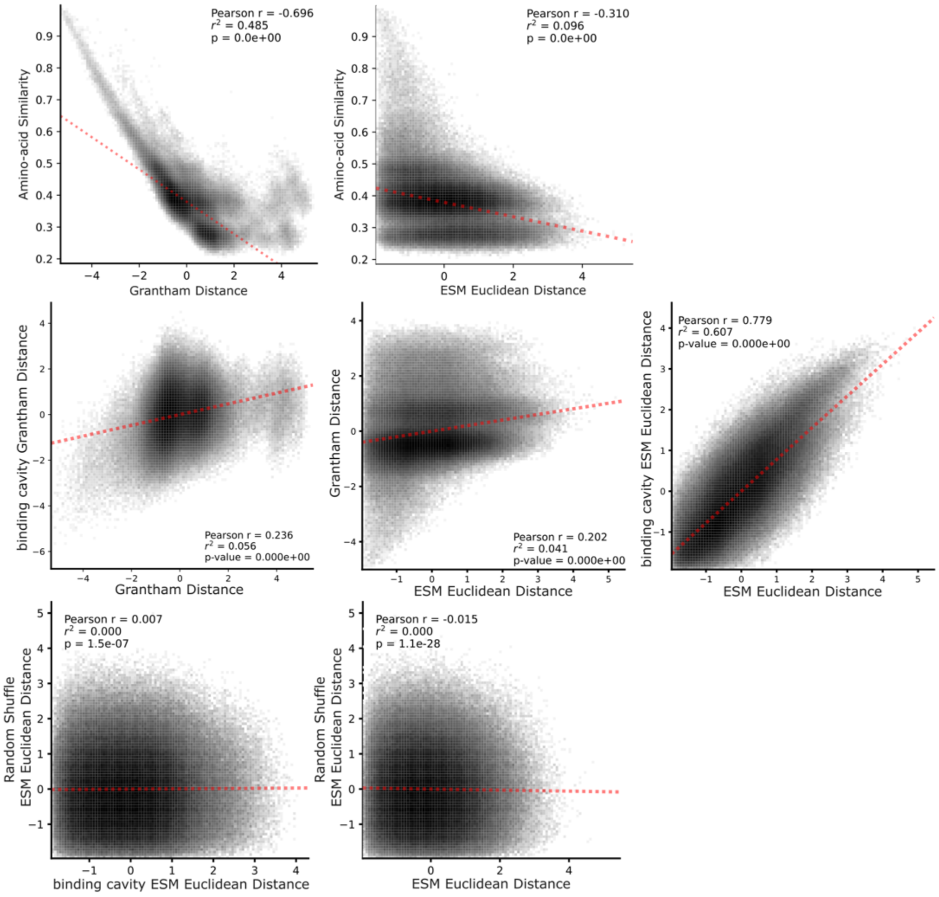


**Figure S2. Comparison of sequence- and embedding-based similarity metrics across olfactory receptors.**

Top: Pearson correlation analysis comparing amino acid similarity–based Grantham distances with ESM-derived Euclidean distances across OR pairs. Middle: Pairwise Pearson correlations among sequence- and structure-informed similarity measures, including full-length sequence Grantham distance vs. binding cavity Grantham distance, Grantham distance vs. ESM Euclidean distance, and binding cavity ESM Euclidean distance vs. full-sequence ESM Euclidean distance. Bottom: Control comparisons against randomized embeddings. Pearson correlations between binding cavity ESM Euclidean distances and randomly shuffled ESM embeddings, and between full-sequence ESM Euclidean distances and shuffled controls, demonstrating preservation of biological signal in learned representations.

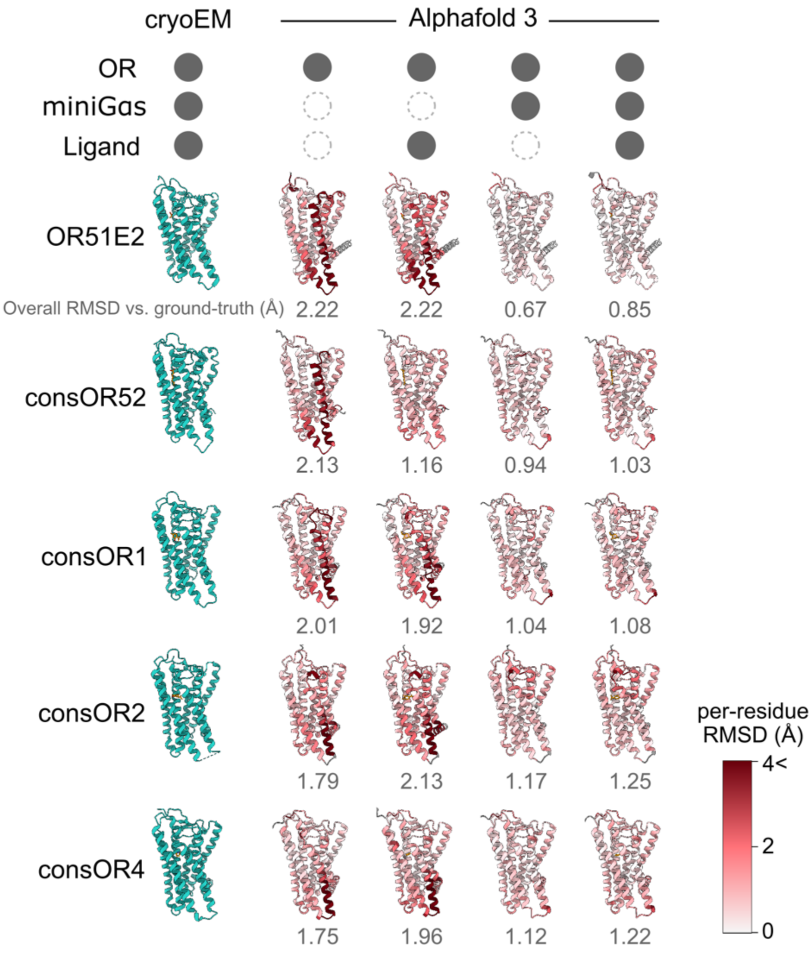


**Figure S3. Structural agreement between cryo-EM olfactory receptor structures and AlphaFold3 predictions.**

Comparison of experimentally determined cryo-EM structures with AlphaFold3-predicted models across multiple modeling conditions, including receptor alone (OR), receptor with ligand (OR + ligand), receptor with mini G protein (OR + miniGαs), and receptor with both ligand and miniGαs protein (OR + ligand + miniGαs). For each receptor, per-residue root-mean-square deviation (RMSD) relative to the cryo-EM structure is shown to assess local structural agreement, alongside overall backbone RMSD values summarized below each model.

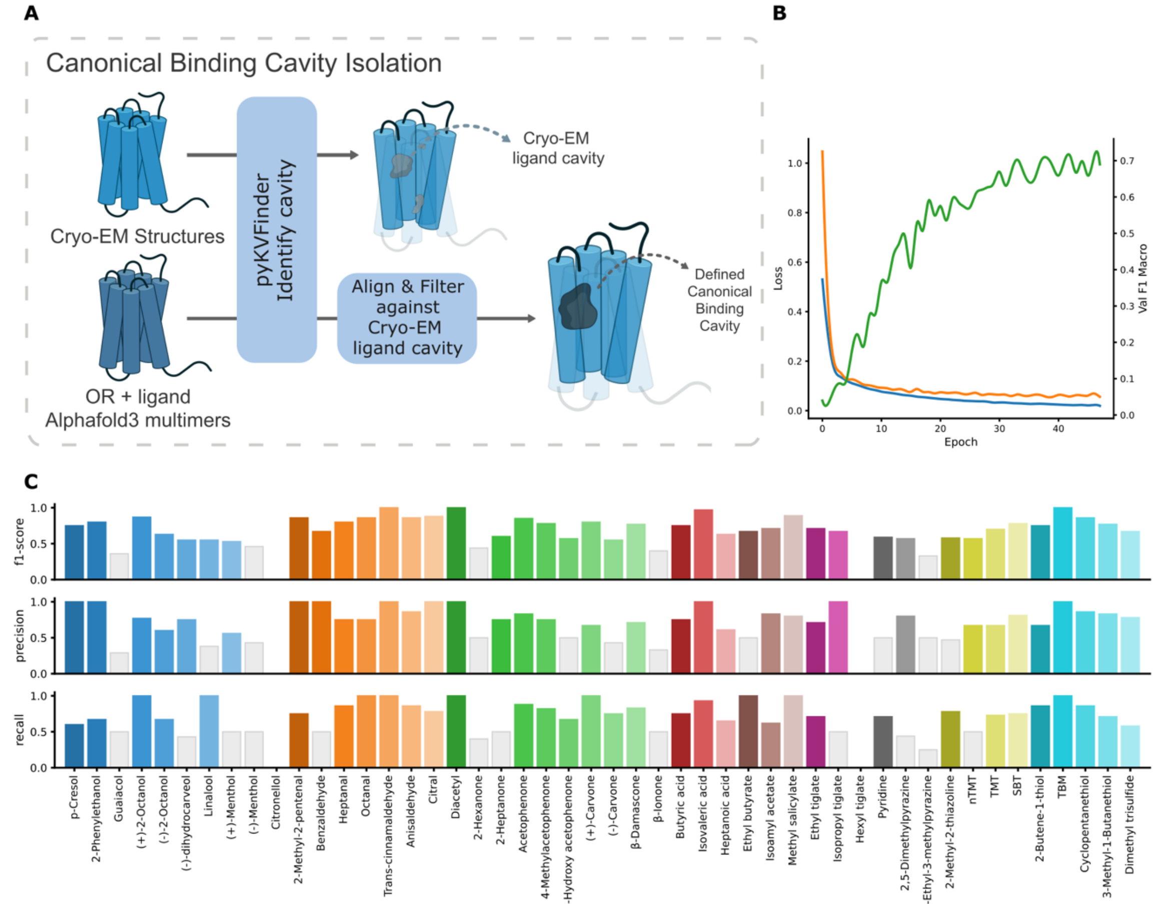


**Figure S4. Binding cavity definition and model performance evaluation.**

**A.** Schematic of the pipeline for canonical binding cavity identification. Cryo-EM structures and corresponding AlphaFold3-predicted models (OR + ligand + miniGαs) were analyzed using pyKVFinder to detect putative cavities. The cryo-EM derived cavity was used as a reference to align and superimpose predicted cavities. The resulting consensus region was defined as the canonical binding cavity used for downstream filtering and voxelization (see Methods for details). **B.** Training dynamics of the final CNN model. Loss curves for training (blue) and validation (orange) are shown across epochs, alongside validation macro F1-score (green), demonstrating stable optimization and concordant performance improvement. **C.** Precision, recall, and F1-score for each of the 48 odorants. Odorants with performance values below 0.5 are highlighted in gray.

### Figure

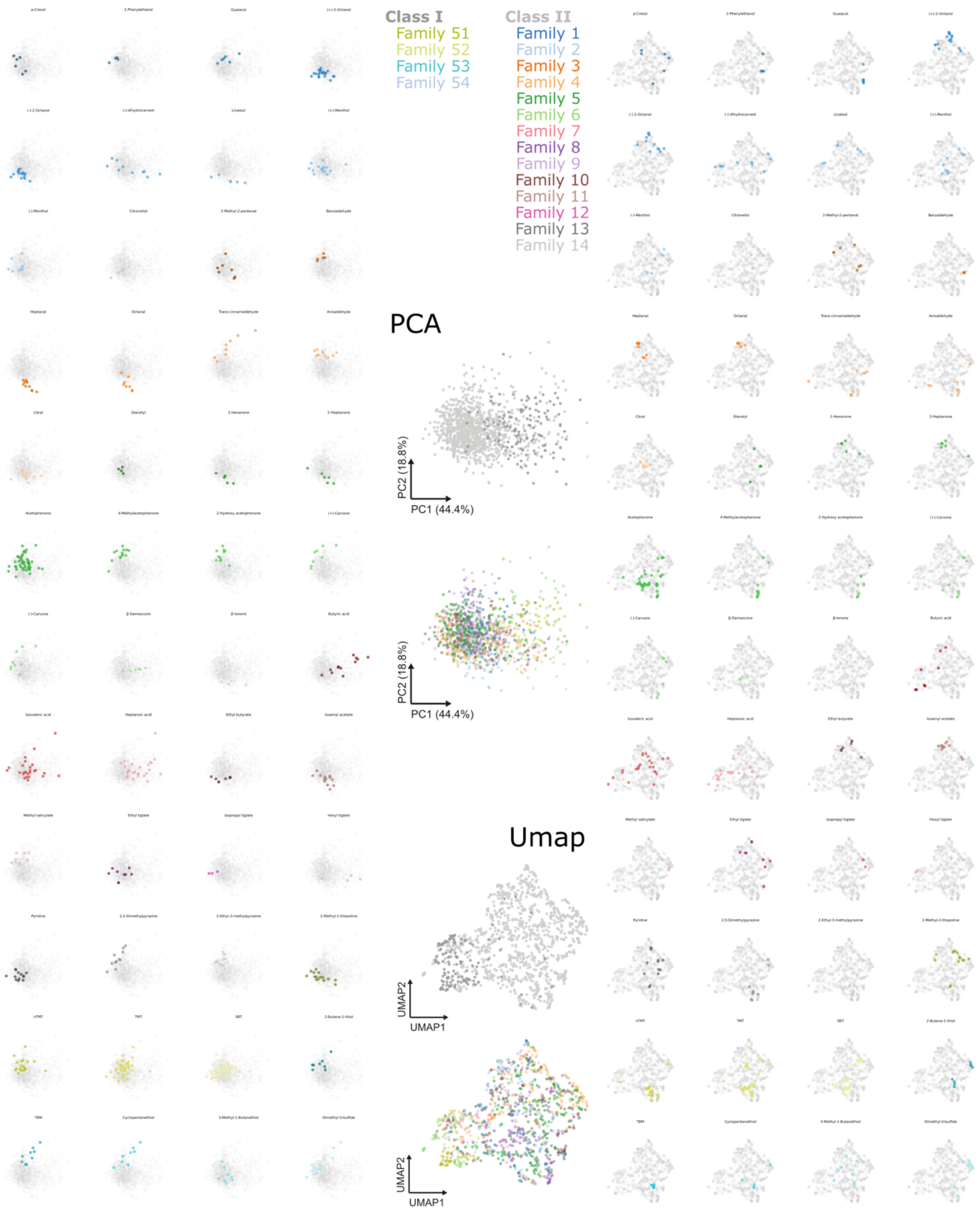
Class I
Family 51
Family 52
Family 53
Family 54
Class II
Family 1
Family 2
Family 3
Family 4
Family 5
Family 6
Family 7
Family 8
Family 9
Family 10
Family 11
Family 12
Family 13
Family 14
PCA
PC2 (18.8%)
PC1 (44.4%)
PC2 (18.8%)
PC1 (44.4%)
Umap
UMAP2
UMAP1
UMAP2
UMAP1

**S5. Global organization of CNN-derived receptor embeddings across dimensionality reduction methods.**

Comprehensive visualization of receptor embeddings using principal component analysis (PCA) and UMAP (smeLLMap). Left: PCA projections showing OR distributions for each of the 48 odorants, with receptors responding to each ligand highlighted. Right: Corresponding UMAP (smeLLMap) projections illustrating odor-specific clustering in the learned embedding space. Middle: Distribution of OR classes and families mapped onto both PCA and UMAP representations, demonstrating that higher-level class separation is preserved while finer family-level organization is less pronounced relative to functional clustering by odor response.

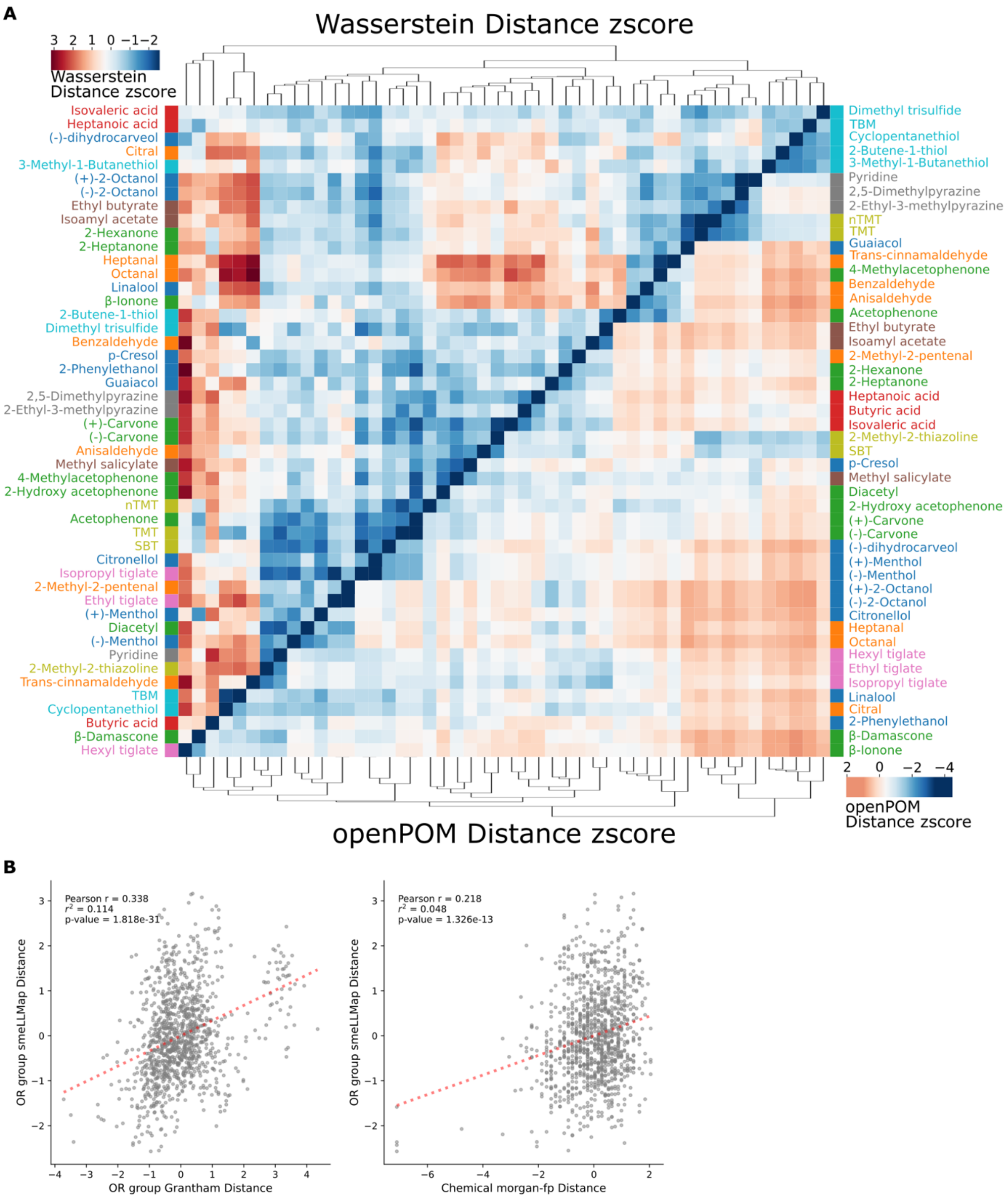
A
Wasserstein Distance zscore
3 2 1 0 −1−2
Wasserstein
Distance zscore
Isovaleric acid
Heptanoic acid
(-)-dihydrocarveol
Citral
3-Methyl-1-Butanethiol
(+)-2-Octanol
(-)-2-Octanol
Ethyl butyrate
Isoamyl acetate
2-Hexanone
2-Heptanone
Heptanal
Octanal
Linalool
β-Ionone
2-Butene-1-thiol
Dimethyl trisulfide
Benzaldehyde
p-Cresol
2-Phenylethanol
Guaiacol
2,5-Dimethylpyrazine
2-Ethyl-3-methylpyrazine
(+)-Carvone
(-)-Carvone
Anisaldehyde
Methyl salicylate
4-Methylacetophenone
2-Hydroxy acetophenone
nTMT
Acetophenone
TMT
SBT
Citronellol
Isopropyl tiglate
2-Methyl-2-pentenal
Ethyl tiglate
(+)-Menthol
Diacetyl
(-)-Menthol
Pyridine
2-Methyl-2-thiazoline
Trans-cinnamaldehyde
TBM
Cyclopentanethiol
Butyric acid
β-Damascone
Hexyl tiglate
Dimethyl trisulfide
TBM
Cyclopentanethiol
2-Butene-1-thiol
3-Methyl-1-Butanethiol
Pyridine
2,5-Dimethylpyrazine
2-Ethyl-3-methylpyrazine
nTMT
TMT
Guaiacol
Trans-cinnamaldehyde
4-Methylacetophenone
Benzaldehyde
Anisaldehyde
Acetophenone
Ethyl butyrate
Isoamyl acetate
2-Methyl-2-pentenal
2-Hexanone
2-Heptanone
Heptanoic acid
Butyric acid
Isovaleric acid
2-Methyl-2-thiazoline
SBT
p-Cresol
Methyl salicylate
Diacetyl
2-Hydroxy acetophenone
(+)-Carvone
(-)-Carvone
(-)-dihydrocarveol
(+)-Menthol
(-)-Menthol
(+)-2-Octanol
(-)-2-Octanol
Citronellol
Heptanal
Octanal
Hexyl tiglate
Ethyl tiglate
Isopropyl tiglate
Linalool
Citral
2-Phenylethanol
β-Damascone
β-Ionone
2 0 −2 −4
openPOM
Distance zscore
openPOM Distance zscore
B
Pearson r = 0.338
$r^2$ = 0.114
p-value = 1.818e-31
OR group smeLLMap Distance
OR group Grantham Distance
Pearson r = 0.218
$r^2$ = 0.048
p-value = 1.326e-13
OR group smeLLMap Distance
Chemical morgan-fp Distance

**Figure S6. Comparison of embedding-derived receptor distances with sequence- and chemistry-based similarity metrics.**

**A.** Dual-clustered heatmaps comparing normalized distance metrics across odorants. Top: z-scored Wasserstein distances between distributions of OR embeddings (smeLLMap) for each odorant pair. Bottom: corresponding z-scored Euclidean distances in OpenPOM space, representing perception-informed chemical similarity. **B.** Correlation analyses comparing receptor embedding distances with alternative similarity measures. Left: Pearson correlation between OR group distances in smeLLMap (Wasserstein) and sequence-based Grantham distances. Right: Pearson correlation between OR group distances in smeLLMap and chemical similarity derived from Morgan fingerprint representations

| | aucroc | prob_corr | f1-score | precision | recall |
|---|---|---|---|---|---|
| p-Cresol | 0.89 | 0.73255 | 0.75 | 1 | 0.6 |
| 2-Phenylethanol | 0.94224422 | 0.80710558 | 0.8 | 1 | 0.67 |
| Guaiacol | 0.97512438 | 0.49389872 | 0.36 | 0.29 | 0.5 |
| (+)-2-Octanol | 1 | 0.91053396 | 0.87 | 0.77 | 1 |
| (-)-2-Octanol | 0.96201814 | 0.67341891 | 0.63 | 0.6 | 0.67 |
| (-)-dihydrocarveol | 0.96392496 | 0.69319504 | 0.55 | 0.75 | 0.43 |
| Linalool | 1 | 0.72351811 | 0.55 | 0.38 | 1 |
| (+)-Menthol | 0.85589744 | 0.62406313 | 0.53 | 0.56 | 0.5 |
| (-)-Menthol | 0.9480737 | 0.5197812 | 0.46 | 0.43 | 0.5 |
| Citronellol | 0.96393035 | 0.37660607 | 0 | 0 | 0 |
| 2-Methyl-2-pentenal | 0.97263682 | 0.84515149 | 0.86 | 1 | 0.75 |
| Benzaldehyde | 0.98275862 | 0.6881775 | 0.67 | 1 | 0.5 |
| Heptanal | 0.98917749 | 0.73052781 | 0.8 | 0.75 | 0.86 |
| Octanal | 1 | 0.90210967 | 0.86 | 0.75 | 1 |
| Trans-cinnamaldehyde | 1 | 0.9859638 | 1 | 1 | 1 |
| Anisaldehyde | 0.9963925 | 0.86971289 | 0.86 | 0.86 | 0.86 |
| Citral | 0.99489796 | 0.86625893 | 0.88 | 1 | 0.78 |
| Diacetyl | 1 | 0.89681083 | 1 | 1 | 1 |
| 2-Hexanone | 0.962 | 0.52969527 | 0.44 | 0.5 | 0.4 |
| 2-Heptanone | 0.99497487 | 0.75804466 | 0.6 | 0.75 | 0.5 |
| Acetophenone | 0.95014094 | 0.81457382 | 0.85 | 0.83 | 0.88 |
| 4-Methylacetophenone | 0.95829428 | 0.83288983 | 0.78 | 0.75 | 0.82 |
| 2-Hydroxy acetophenone | 0.96901173 | 0.62587029 | 0.57 | 0.5 | 0.67 |
| (+)-Carvone | 1 | 0.90482121 | 0.8 | 0.67 | 1 |
| (-)-Carvone | 0.99129353 | 0.70908125 | 0.55 | 0.43 | 0.75 |
| Œ≤-Damascone | 0.9120603 | 0.86859694 | 0.77 | 0.71 | 0.83 |
| Œ≤-Ionone | 0.94581281 | 0.49842657 | 0.4 | 0.33 | 0.5 |
| Butyric acid | 0.99875622 | 0.88191084 | 0.75 | 0.75 | 0.75 |
| Isovaleric acid | 0.99719298 | 0.91817394 | 0.97 | 1 | 0.93 |
| Heptanoic acid | 0.89893617 | 0.6611081 | 0.63 | 0.61 | 0.65 |
| Ethyl butyrate | 0.99261084 | 0.57559229 | 0.67 | 0.5 | 1 |
| Isoamyl acetate | 0.95748731 | 0.74228959 | 0.71 | 0.83 | 0.62 |
| Methyl salicylate | 1 | 0.90854191 | 0.89 | 0.8 | 1 |
| Ethyl tiglate | 0.98556999 | 0.71354159 | 0.71 | 0.71 | 0.71 |

| | | | | | |
|---|---|---|---|---|---|
| Isopropyl tiglate | 1 | 0.79596373 | 0.67 | 1 | 0.5 |
| Hexyl tiglate | 0.9141791 | 0.05535212 | 0 | 0 | 0 |
| Pyridine | 0.85930736 | 0.67125846 | 0.59 | 0.5 | 0.71 |
| 2,5-Dimethylpyrazine | 0.9739229 | 0.67666771 | 0.57 | 0.8 | 0.44 |
| 2-Ethyl-3-methylpyrazine | 0.93159204 | 0.38868748 | 0.33 | 0.5 | 0.25 |
| 2-Methyl-2-thiazoline | 0.93877551 | 0.63984346 | 0.58 | 0.47 | 0.78 |
| nTMT | 0.75124378 | 0.64863775 | 0.57 | 0.67 | 0.5 |
| TMT | 0.96969697 | 0.76199443 | 0.7 | 0.67 | 0.73 |
| SBT | 0.94612591 | 0.75225294 | 0.78 | 0.81 | 0.75 |
| 2-Butene-1-thiol | 0.86075036 | 0.7815141 | 0.75 | 0.67 | 0.86 |
| TBM | 1 | 0.99962926 | 1 | 1 | 1 |
| Cyclopentanethiol | 0.998557 | 0.9202946 | 0.86 | 0.86 | 0.86 |
| 3-Methyl-1-Butanethiol | 0.98124098 | 0.80131139 | 0.77 | 0.83 | 0.71 |
| Dimethyl trisulfide | 0.98272884 | 0.68576128 | 0.67 | 0.78 | 0.58 |

**Table S1. Trained model's validation odor prediction metric**

Summary of validation performance for each odorant, including area under the receiver operating characteristic curve (AUROC), probability–response correlation (prob_corr), F1-score, precision, and recall.

| | odor | cid | n_OR | obs_grantham_mean | obs_grantham_median | null_grantham_mean | null_grantham_median | p_grantham_perm_mean | p_grantham_perm_median | obs_euclid_mean | obs_euclid_median | null_euclid_mean | null_euclid_median | p_euclid_perm_mean | p_euclid_perm_median |
|---|---|---|---|---|---|---|---|---|---|---|---|---|---|---|---|
| 0 | Dimethyl trisulfide | 19310 | 16 | 0.0139 | 0.2510 | 0.0045 | 0.0374 | 0.5020 | 0.8260 | -0.4434 | -0.9146 | 0.0121 | -0.1635 | 0.0560 | 0.0000 |
| 1 | 3-Methyl-1-Butanethiol | 10925 | 8 | -0.6002 | -0.6424 | -0.0052 | 0.0222 | 0.0360 | 0.0320 | -0.5814 | -0.5565 | -0.0139 | -0.1509 | 0.0600 | 0.1320 |
| 2 | Cyclopentanethiol | 15510 | 9 | -1.2545 | -0.7162 | 0.0070 | 0.0321 | 0.0000 | 0.0080 | -0.2826 | -0.4438 | 0.0395 | -0.1076 | 0.1980 | 0.1780 |
| 3 | TBM | 6387 | 10 | -1.3706 | -0.8363 | -0.0123 | 0.0275 | 0.0000 | 0.0020 | -0.7029 | -0.7895 | 0.0199 | -0.1409 | 0.0180 | 0.0140 |
| 4 | 2-Butene-1-thiol | 6433451 | 9 | -0.1284 | -0.0562 | -0.0218 | 0.0145 | 0.3740 | 0.4240 | -0.7640 | -0.7688 | 0.0242 | -0.1264 | 0.0160 | 0.0280 |
| 5 | SBT | 162148 | 49 | -0.3311 | -0.2973 | -0.0039 | 0.0218 | 0.0000 | 0.0000 | -0.7523 | -0.8661 | 0.0069 | -0.1583 | 0.0000 | 0.0000 |
| 6 | TMT | 263626 | 45 | -0.3582 | -0.2838 | -0.0032 | 0.0135 | 0.0020 | 0.0140 | -0.6614 | -0.8130 | 0.0141 | -0.1579 | 0.0000 | 0.0000 |
| 7 | nTMT | 61653 | 19 | -0.6101 | -0.6017 | -0.0061 | -0.0018 | 0.0000 | 0.0000 | -0.5728 | -0.6272 | 0.0089 | -0.1558 | 0.0140 | 0.0240 |
| 8 | 2-Methyl-2-thiazoline | 16867 | 15 | 0.1310 | 0.1996 | 0.0058 | 0.0274 | 0.7020 | 0.7780 | -0.7255 | -0.9066 | -0.0142 | -0.1840 | 0.0040 | 0.0000 |
| 9 | 2-Ethyl-3-methylpyrazine | 27457 | 9 | -0.1909 | 0.1550 | -0.0033 | 0.0011 | 0.2520 | 0.6840 | -0.9526 | -0.8981 | 0.0094 | -0.1229 | 0.0060 | 0.0160 |
| 10 | 2,5-Dimethylpyrazine | 31252 | 11 | -1.0211 | -0.7582 | 0.0051 | 0.0426 | 0.0000 | 0.0040 | -0.6545 | -0.7139 | 0.0001 | -0.1873 | 0.0160 | 0.0220 |
| 11 | Pyridine | 1049 | 13 | -0.1694 | -0.0762 | 0.0033 | 0.0198 | 0.2280 | 0.3220 | -0.9442 | -1.0281 | 0.0188 | -0.1384 | 0.0000 | 0.0000 |
| 12 | Hexyl tiglate | 637523 | 4 | 1.1755 | 1.0706 | 0.0027 | 0.0449 | 0.9920 | 0.9740 | 0.0642 | -0.0640 | 0.0033 | -0.0989 | 0.5840 | 0.5220 |
| 13 | Isopropyl tiglate | 5367745 | 3 | -0.8005 | -0.5705 | 0.0188 | 0.0704 | 0.1140 | 0.1960 | -1.4211 | -1.4639 | | | 0.0120 | 0.0100 |
| 14 | Ethyl tiglate | 5281163 | 10 | -0.2343 | 0.1063 | -0.0070 | 0.0057 | 0.2480 | 0.6020 | -0.5717 | -0.5491 | -0.0331 | -0.1650 | 0.0800 | 0.1520 |
| 15 | Methyl salicylate | 4133 | 9 | -0.5034 | -0.4769 | -0.0134 | -0.0001 | 0.0360 | 0.0420 | -1.0183 | -1.0613 | 0.0065 | -0.1460 | 0.0020 | 0.0020 |
| 16 | Isoamyl acetate | 31276 | 15 | -0.6118 | -0.7165 | -0.0032 | 0.0290 | 0.0000 | 0.0000 | -0.9296 | -0.9819 | 0.0114 | -0.1349 | 0.0000 | 0.0000 |
| 17 | Ethyl butyrate | 7762 | 9 | -1.0085 | -0.4191 | 0.0045 | 0.0060 | 0.0000 | 0.0860 | -0.9256 | -0.5746 | -0.0030 | -0.1478 | 0.0040 | 0.0900 |
| 18 | Heptanoic acid | 8094 | 29 | -0.0594 | -0.0132 | 0.0006 | 0.0212 | 0.3500 | 0.4100 | -0.1697 | -0.3239 | 0.0147 | -0.1579 | 0.1720 | 0.1860 |
| 19 | Isovaleric acid | 10430 | 25 | 0.0693 | 0.0724 | -0.0045 | 0.0086 | 0.6660 | 0.6380 | -0.1424 | -0.4405 | -0.0009 | -0.1642 | 0.2600 | 0.0780 |
| 20 | Butyric acid | 264 | 13 | -0.8434 | -0.5156 | 0.0001 | 0.0394 | 0.0000 | 0.0140 | 0.0322 | -0.1850 | 0.0103 | -0.1571 | 0.5200 | 0.4680 |
| 21 | Œ≤-Ionone | 638014 | 4 | -0.2785 | -0.3560 | 0.0469 | 0.1100 | 0.2640 | 0.2260 | -0.4353 | -0.6930 | 0.0021 | -0.0907 | 0.2560 | 0.1220 |
| 22 | Œ≤-Damascone | 32052 | 6 | -0.2710 | -0.1163 | -0.0173 | 0.0429 | 0.2620 | 0.3660 | -0.8630 | -0.8647 | -0.0101 | -0.1489 | 0.0320 | 0.0360 |
| 23 | (-)-Carvone | 439570 | 9 | -0.9101 | -0.6223 | 0.0132 | 0.0439 | 0.0020 | 0.0220 | -0.2588 | -0.3203 | 0.0115 | -0.1372 | 0.2560 | 0.3120 |
| 24 | (+)-Carvone | 16724 | 8 | -1.0134 | -0.8159 | -0.0174 | 0.0083 | 0.0040 | 0.0160 | -0.3974 | -0.6011 | -0.0206 | -0.1714 | 0.1900 | 0.1200 |
| 25 | 2-Hydroxyacetophenone | 68490 | 11 | -1.2471 | -0.7958 | -0.0041 | 0.0402 | 0.0000 | 0.0060 | -1.0496 | -1.1636 | -0.0162 | -0.1591 | 0.0000 | 0.0000 |
| 26 | 4-Methylacetophenone | 8500 | 14 | -1.5150 | -1.1524 | -0.0035 | 0.0306 | 0.0000 | 0.0000 | -0.9671 | -1.0855 | 0.0329 | -0.1243 | 0.0000 | 0.0000 |
| 27 | Acetophenone | 7410 | 50 | -0.4086 | -0.3580 | -0.0056 | 0.0189 | 0.0000 | 0.0000 | -0.6622 | -0.7055 | -0.0113 | -0.1744 | 0.0000 | 0.0000 |
| 28 | 2-Heptanone | 8051 | 6 | -1.0715 | -1.0579 | -0.0062 | 0.0150 | 0.0060 | 0.0060 | -0.7345 | -1.0901 | 0.0101 | -0.1677 | 0.0480 | 0.0040 |
| 29 | 2-Hexanone | 11583 | 5 | -1.5064 | -1.6568 | 0.0125 | 0.0608 | 0.0000 | 0.0000 | -0.9709 | -0.9581 | 0.0597 | -0.0765 | 0.0120 | 0.0200 |
| 30 | Diacetyl | 650 | 4 | -0.2798 | -0.1759 | -0.0133 | 0.0248 | 0.3340 | 0.3820 | -1.4941 | -1.5429 | -0.0021 | -0.0714 | 0.0000 | 0.0000 |
| 31 | Citral | 638011 | 12 | -0.8474 | -0.8365 | -0.0069 | 0.0208 | 0.0000 | 0.0000 | -0.9139 | -1.0209 | -0.0101 | -0.1720 | 0.0000 | 0.0000 |
| 32 | Anisaldehyde | 31244 | 11 | -0.7312 | -0.2821 | -0.0174 | 0.0070 | 0.0100 | 0.1440 | -1.0188 | -1.0694 | 0.0347 | -0.1174 | 0.0020 | 0.0000 |

| | | | | | | | | | | | | | | | |
|---|---|---|---|---|---|---|---|---|---|---|---|---|---|---|---|
| 33 | Trans-cinnamaldehyde | 637511 | 8 | 0.1378 | 0.2113 | 0.0124 | 0.0123 | 0.6700 | 0.6960 | 0.1259 | -0.1036 | 0.0008 | -0.1271 | 0.6420 | 0.5280 |
| 34 | Octanal | 454 | 8 | -1.6014 | -1.3522 | -0.0002 | 0.0241 | 0.0000 | 0.0000 | -1.1745 | -1.0998 | -0.0159 | -0.1651 | 0.0000 | 0.0020 |
| 35 | Heptanal | 8130 | 11 | -0.9677 | -0.8765 | 0.0087 | 0.0237 | 0.0000 | 0.0020 | -1.2894 | -1.3305 | 0.0007 | -0.1670 | 0.0000 | 0.0000 |
| 36 | Benzaldehyde | 240 | 5 | 0.2295 | -0.0425 | 0.0084 | 0.0230 | 0.6980 | 0.4600 | -1.4918 | -1.3870 | 0.0216 | -0.1232 | 0.0000 | 0.0000 |
| 37 | 2-Methyl-2-pentenal | 5319754 | 7 | -0.4334 | -0.0888 | -0.0015 | -0.0007 | 0.1000 | 0.3900 | -0.4622 | -0.4358 | 0.0059 | -0.1330 | 0.1480 | 0.2500 |
| 38 | Citronellol | 8842 | 5 | -0.4148 | 0.1307 | -0.0095 | 0.0116 | 0.1720 | 0.6000 | -1.1600 | -1.3094 | -0.0015 | -0.1350 | 0.0060 | 0.0000 |
| 39 | (-)-Menthol | 16666 | 9 | -0.5812 | -0.6679 | -0.0212 | 0.0061 | 0.0400 | 0.0240 | -0.9245 | -0.9452 | -0.0225 | -0.1713 | 0.0060 | 0.0120 |
| 40 | (+)-Menthol | 165675 | 15 | -0.9349 | -0.8441 | -0.0050 | 0.0167 | 0.0000 | 0.0000 | -0.6592 | -0.7303 | 0.0084 | -0.1751 | 0.0020 | 0.0060 |
| 41 | Linalool | 6549 | 6 | -0.9286 | -0.8246 | -0.0060 | 0.0142 | 0.0120 | 0.0380 | -0.4534 | -0.3529 | 0.0261 | -0.0989 | 0.1460 | 0.2800 |
| 42 | (-)-dihydrocarveol | 443163 | 10 | -0.2008 | -0.1261 | 0.0156 | 0.0478 | 0.2320 | 0.3160 | 0.5272 | 0.2167 | -0.0021 | -0.1701 | 0.9200 | 0.8340 |
| 43 | (-)-2-Octanol | 80080 | 17 | -0.8222 | -0.8550 | -0.0043 | 0.0277 | 0.0000 | 0.0000 | -1.1240 | -1.1137 | 0.0057 | -0.1652 | 0.0000 | 0.0000 |
| 44 | (+)-2-Octanol | 2723888 | 22 | -0.2801 | -0.3046 | 0.0000 | 0.0317 | 0.0680 | 0.0640 | -0.9740 | -1.0792 | 0.0128 | -0.1573 | 0.0000 | 0.0000 |
| 45 | Guaiacol | 460 | 6 | -0.8643 | -0.3343 | -0.0144 | 0.0206 | 0.0140 | 0.1860 | -1.0240 | -1.2228 | -0.0112 | -0.1419 | 0.0040 | 0.0020 |
| 46 | 2-Phenylethanol | 6054 | 4 | -1.0953 | -0.9677 | -0.0570 | 0.0070 | 0.0200 | 0.0420 | -1.2577 | -1.3415 | 0.0196 | -0.0261 | 0.0020 | 0.0000 |
| 47 | p-Cresol | 2879 | 6 | -0.8059 | -0.7866 | -0.0238 | 0.0237 | 0.0260 | 0.0480 | -0.4143 | -0.3852 | -0.0011 | -0.1114 | 0.2060 | 0.2760 |

**Table S2. Pairwise receptor similarity metrics for odor-specific responding ORs.**

Summary of observed and permuted pairwise distances among ORs responding to each odorant, computed using Grantham sequence distances and smeLLMap embedding Euclidean distances. For each odorant, mean and median distances are reported alongside corresponding null distributions generated by random permutation, with associated permutation-based *p-values* indicating statistical significance.

7. A. Ravia *et al.*, A measure of smell enables the creation of olfactory metamers. *Nature* **588**, 118-123 (2020).
8. H. L. Morgan, The generation of a unique machine description for chemical structures-a technique developed at chemical abstracts service. *Journal of chemical documentation* **5**, 107-113 (1965).
9. D. Rogers, M. Hahn, Extended-connectivity fingerprints. *Journal of chemical information and modeling* **50**, 742-754 (2010).
10. B. K. Lee *et al.*, A principal odor map unifies diverse tasks in olfactory perception. *Science* **381**, 999-1006 (2023).
11. A. A. Barsainyan, R. Kumar, P. Saha, M. Schmuker. (2023).
12. A. Keller, L. B. Vosshall, Olfactory perception of chemically diverse molecules. *BMC Neurosci* **17**, 55 (2016).
13. K. Katoh, K.-i. Kuma, H. Toh, T. Miyata, MAFFT version 5: improvement in accuracy of multiple sequence alignment. *Nucleic acids research* **33**, 511-518 (2005).
14. D. Darriba *et al.*, ModelTest-NG: a new and scalable tool for the selection of DNA and protein evolutionary models. *Molecular biology and evolution* **37**, 291-294 (2020).
15. R. Grantham, Amino acid difference formula to help explain protein evolution. *science* **185**, 862-864 (1974).
16. J. Abramson *et al.*, Accurate structure prediction of biomolecular interactions with AlphaFold 3. *Nature* **630**, 493-500 (2024).
17. Y. Zhang, J. Skolnick, TM-align: a protein structure alignment algorithm based on the TM-score. *Nucleic acids research* **33**, 2302-2309 (2005).
18. J. V. d. S. Guerra *et al.*, pyKVFinder: an efficient and integrable Python package for biomolecular cavity detection and characterization in data science. *BMC Bioinformatics* **22**, 607 (2021).
19. H. Saito, M. Kubota, R. W. Roberts, Q. Chi, H. Matsunami, RTP family members induce functional expression of mammalian odorant receptors. *Cell* **119**, 679-691 (2004).
20. H. Zhuang, H. Matsunami, Evaluating cell-surface expression and measuring activation of mammalian odorant receptors in heterologous cells. *Nat Protoc* **3**, 1402-1413 (2008).
